# Astaxanthin extends lifespan via altered biogenesis of the mitochondrial respiratory chain complex III

**DOI:** 10.1101/698001

**Authors:** Ronit Hoffman, Laure D. Sultan, Ann Saada, Joseph Hirschberg, Oren Osterzetser-Biran, Yosef Gruenbaum

## Abstract

Astaxanthin is a *keto*-carotenoid produced in some bacteria and algae, which has very important industrial applications (i.e., in cosmetics, coloring additive in aquaculture and as a dietary supplement for human). Here, we analyzed the molecular basis of Astaxanthin-mediated prolongevity in the model organism, *Caenorhabditis elegans*. The increased lifespan effects of Astaxanthin are restricted in *C. elegans* to the adult phase and are uninfluenced by various other carotenoids tested. Genetic analyses indicated that the Astaxanthin-mediated life-extension relies on mitochondria activity, via the Rieske iron-sulfur polypeptide-1 (ISP-1), but is not influenced by the functions of other known longevity-related gene-loci, including *CLK-1, DAF-2*, *DAT-16*, *EAT-2*, *GAS-1 GLP-1* or *MEV-1*. Biochemical analyses of native respiratory complexes showed that Astaxanthin affects the biogenesis of holo-complex III (and likely supercomplex I+III, as well). Effects on holo-CIII assembly and activity were also indicated by *in-vitro* assays, with mitochondria isolated from worms, rodents, human and plants, which were treated with Astaxanthin. These data indicated a cross-species effect on the oxidative phosphorylation (OXPHOS) machinery by the carotenoid, and provide with further insights into the molecular mechanism of animals longevity extension by Astaxanthin.

**Significance Statement:** Astaxanthin is a widely consumed pigment by animals and human. In this study we find that Astaxanthin, but not other tested carotenoids, significantly extends the lifespan of animals by affecting respiratory complex III (CIII) biogenesis of the mitochondria, in plants, C. elegans, rodents and human. We further propose a model to try explaining this effect of astaxanthin on animals’ longevity.

## Introduction

The nematode *Caenorhabditis elegans* is a leading model organism for studying the biology of aging, and for identifying new pharmacological targets for extended longevity and the treatment of aging-related diseases in animals (Chen *et al*. 2014). Genetic studies indicated to three major aging-related mechanisms in *C. elegans*. These include the insulin pathway, the dietary restriction system, and the mitochondrial respiration system, which were all shown to affect the lifespan of worms and other animals (Amrit *et al*. 2014, Collins *et al*. 2008). Noteworthy, the majority (>80%) of the *C. elegans* proteome was shown to share homology with known human genes (Henricson *et al*. 2004, Lai *et al*. 2000), thus suggesting common aging-related mechanisms between *C. elegans* and human.

Among the key factors that affect longevity in worms is oxidative damage, generated by <u>r</u>eactive <u>o</u>xygen <u>s</u>pecies (ROS). Induced ROS level was shown to induce early aging in various animals and plants (Paital *et al*. 2016, Singh *et al*. 2016). Consequently, strong antioxidants as resveratrol, coenzyme Q10 (ubiquinol or 2,3-dimethoxy-5-methyl-6-decaprenyl benzoquinone) and carotenoids as Astaxanthin can extend the lifespan of *C. elegans*, presumably by scavenging ROS and lowering the levels of oxidative stress (Chen*, et al.* 2014). Carotenoids comprise a large class of natural antioxidant pigments that are synthetized and accumulate in photosynthetic organisms, various bacteria and some fungi (Cazzonelli 2011, Rodriguez-Concepcion et al. 2018). These isoprenoid compounds are also essential components of the photosynthetic machinery, and are associated with anti-oxidative reactions in bacteria, fungi, algae and plants. In animals, carotenoids that are consumed from the food play important roles in the antioxidant defense mechanism (Fiedor and Burda 2014).

Astaxanthin (i.e., 3,3-dihydroxy-β,β-carotene-4,4-dione) is an orange-red pigment containing 13 conjugated double bonds (Supplementary Fig. S1). As other carotenoids, Astaxanthin is a lipid-soluble pigment, which has self-limited absorption orally, with no known toxic syndromes (Barros *et al*. 2014). Astaxanthin is naturally produced by some marine bacteria, algae and plants. Many red-colored marine animals (e.g., salmons, shrimps) and a few bird species (e.g., flamingo) obtain the Astaxanthin orally, through the food chain (Ambati *et al*. 2014). As a strong antioxidant, Astaxanthin was utilized as a common antioxidant agent and as a dietary supplement, intended for human, animal and aquaculture consumption. Astaxanthin has been implicated in numerous health benefits in humans, including reduction in cardiovascular diseases, enhancement of the immune response and reduction in the occurrence of various cancers. (Ames 2018, Zhang and Wang 2015, Zhang *et al*. 2016). Astaxanthin is obtained industrially by either chemical synthesis or by extraction from algae or bacteria, both of which are approved by the U.S. Food and Drug Administration as food, feed or coloring additives (Raposo *et al*. 2015, Zhang and Wang 2015). Plant-derived Astaxanthin is generally recognized as safe by the FDA, meaning it can be sold as a dietary supplement (Zhang and Wang 2015), although in the United States is restricted to be used only as a colorant agent to animal feed. Currently, there is great interest around Astaxanthin due to its biochemical characteristics, mainly as a potent antioxidant, which is approximately 10 times more effective than β-carotene or lutein and about 100 times than α-tocopherol (Higuera-Ciapara *et al*. 2006, Rao *et al*. 2015). These studies further suggest that Astaxanthin affects cellular metabolism and protects against fatty acid oxidation.

Astaxanthin was previously shown to extent the lifespan of both flies and worms. *Drosophila melanogaster* mutants in CuZn-superoxide dismutase (SOD1) and Mn-superoxide dismutase (SOD2), as well as mutant-lines with reduced levels of SOD1, SOD2 and catalase, show significant lifespan extension and amelioration of age-related decline in motility when are fed with *Haematococcus pluvialis* (a microalga which accumulates high-levels of Astaxanthin) (Huangfu *et al*. 2013). Likewise, feeding *C. elegans* with Astaxanthin also extends the mean lifespans of both wild-type and long-lived mutant *age-1* animals (Kashima et al. 2012, Yazaki et al. 2011). The molecular basis of astaxanthin-mediated prolongevity effects seen in both *D. melanogaster* and *C. elegans* remained unknown, but it was anticipated to be mediated through the scavenging of reactive oxygen species (ROS) produced in these animals by the carotenoid compound.

In this study we show that the lifespan of *C. elegans* fed with Astaxanthin increases by about 20%, which is in accordance with previously publish data showing the effects of Astaxanthin on the mean lifespans of the warms (Liu *et al*. 2016). Our data further indicate that the effects of Astaxanthin are independent of the insulin/IGF or dietary restriction of germline pathways, and provide with evidence that the extended longevity effects on *C. elegans* is mediated by the disassembly or altered biogenesis of the native respiratory complexes III_2_ and presumably the suppercomplex I+III_2_. The effects of Astaxanthin on the mitochondrial OXPHOS system seems universal, as was also apparent by ‘*in-organello*’ assays with mitochondria preparations from plant, *C. elegans*, mice rat and human. These results are of great importance as they offer with novel insights into the molecular basis of longevity extension by Astaxanthin, and may also lead to developing new generation of drugs based on carotenoids to prolong lifespan in animals and human.

## Materials and Methods

### C. elegans strains

*C. elegans* strains maintenance was performed under standard conditions, essentially as described (Brenner 1974). N2 (Wild-type), *daf-2*(e1370), *daf-16*(mu86), *glp-1*(or178), *eat-2*(ad1116), *clk-1*(e2519), *mev-1*(kn1), *gas-1*(fc21) and *isp-1*(qm150) strains were used in this study. All strains were obtained from the *C. elegans* Genome Center (CGC) and were out-crossed with wild-type (N2) animals.

### Lifespan assays

Lifespan assays were performed essentially as described previously (Bar *et al*. 2016). Synchronized *C. elegans* animals were grown on NGM (K_2_HPO_4_, KH_2_PO_4,_ NaCl, Bacto peptone, Agar, Agarose, MgSO_4_, CaCl_2_, and Cholesterol) plates containing one of the following bacterial strains: Wild type *Paracoccus marcusii,* which are accumulating high-levels of Astaxanthin, *P. marcusii* ΔcrtB mutant which lacks carotenoids, *Escherichia coli* OP50 strain (Taxonomy ID 637912), recombinant *E. coli* cells carrying the pASTA plasmid for the expression of Astaxanthin (Cunningham and Gantt 2007). The carotenoid composition in different bacterial strains used in this work is indicated in Supplementary Table S1. In addition, the worms were also grown in the presence of *E. coli* cells in agar-plates supplemented with Astaxanthin (extracted from the algae *Haematococcus pluvialis*) solubilized in 5% DMSO (w/v), and agar-plates containing 5% DMSO with *E. coli* (OP50) cells, as control. Experiments were performed at 20°C, with the exception of *glp-1* and *daf-2* mutant-lines which were conducted at 25°C. In the *daf-2* experiments, the worms were grown at 16°C until the L4 stage, and then transferred to 25°C. For each experiment, 80 adult animals were placed on four separate NGM-plates. Animals were considered dead when they no longer responded to gentle prodding with a platinum wire. Scoring was performed every day. For all lifespan experiments, assays were repeated at least twice.

### Measurements of carotenoids concentration

Wild-type *P. marcusii* or *E. coli* producing Astaxanthin were plated on NGM media, harvested from the plates and resuspended in 2 ml of DDW. The cells were then washed once in 2 ml DDW. The bacterial suspension (1.8 ml) was incubated at 105°C for 24 h, after which the weight of the bacteria was measured. For carotenoid extraction, after washing in DDW, the cells were resuspended in 1 ml of acetone and incubated at 65°C for 10 min in the dark. The samples were centrifuged again at 13,000 xg for 10 min and the acetone supernatant, containing the pigments, was placed in a clean tube. The pigment extract was dried with N_2_ and then stored in the freezer at -20°C. Acetone-dissolved pigment extracts were analyzed by high performance liquid chromatography (HPLC), essentially as previously described (Neuman et al. 2014, Ronen et al. 1999), using a Waters 996 photodiode array detector. Carotenoids were identified based on their characteristic absorption spectra and typical retention times, which corresponded to standard compounds (Isaacson *et al*. 2002). For Astaxanthin extraction, wild-type algae *Haematococcus pluvialis* (kindly provided by Prof. Sammy Boussiba, BGU) were grinded with a mortar and pestle in 5 ml of acetone. The cells were incubated at 65°C for 10 min in the dark, and centrifuged at 13,000 xg for 10 min. The supernatant, containing acetone-soluble pigments, was placed in a clean tube, dried with a stream of N_2_ and then subjected to HPLC analysis.

### Crude mitochondrial preparations from *C. elegans* cells

Mitochondria were isolated from *C. elegans* according to the method described in Grad *et al*. (2007). Bleached embryos were grown at 20°C, on NGM plates seeded with either *P. marcusii* expressing Astaxanthin or *E. coli* (OP50). Before extraction, worms of the specified stages were washed off plates with sterile M9 solution (42 mM Na_2_HPO_4_, 22 mM KH_2_PO_4_, 86 mM NaCl, and 1 mM MgSO_4_·7H_2_O). The worms were pelleted by centrifuging at 3,000g for 5 min, and then resuspended in 10 ml of ice-cold isolation buffer (IB) (210 mM mannitol, 70 mM sucrose, 0.1 mM EDTA, 5 mM Tris-HCl pH 7.4) in the presence of protease inhibitors. The worms were homogenized using a glass homogenizer for 15 strokes. The homogenate was collected in a 50 ml tube and 10 ml of IB was added. Then tubes were centrifuged at 750 g for 10 min at 4°C. The clear supernatant was transferred to a new tube, and centrifuged at 12,000 g for 10 min at 4°C to pellet the mitochondria. The organellar fraction was washed once in 10 ml ice-cold IB (12,000g for 10 min at 4°C), and the mitochondrial pellet was aliquoted and stored at -80°C.

### Mitochondrial extraction from cauliflower

Mitochondria were isolated from cauliflower (*Brassica oleracea* var. botrytis) inflorescences, which allows the purification of large quantities of highly enriched organellar preparations (Keren *et al*. 2009, Neuwirt *et al*. 2005, Sultan *et al*. 2016). About 1.0 kg fresh weight inflorescences were cut off from the stems and kept overnight (about 16 h) in cold water (4°C). In the following morning, the inflorescences were ground with ice-cold extraction buffer (0.9 M mannitol, 90 mM Na-pyrophosphate, 6 mM EDTA, 2.4% PVP25 (w/v), 0.9% BSA (w/v), 9 mM cysteine, 15 mM glycine, and 6 mM β-mercaptoethanol; pH 7.5). Mitochondria were recovered from the extract by differential centrifugations and purifisd on Percoll gradients. Following fractionation, the mitochondria were pelleted, resuspended in a small volume of wash buffer (0.3 M mannitol, 10 mM K-phosphate, 1 mM EDTA, pH 7.5), aliquoted and stored frozen at -80°C. For the isolation of mitochondria from Arabidopsis, we used 2-week-old seedlings grown on MS-plates. The enriched organelle preparation was performed essentially as described in (Keren *et al*. 2012). For analysis of Astaxanthin effects, the mitochondria were incubated with Astaxanthin dissolved in DMSO or DMSO alone for 5 min on ice.

### Mitochondrial enriched preparation from mammals

Mitochondrial enriched fraction, was prepared from human (fibroblast) and mouse (heart) by teflon-glass homogenization and differential centrifugation in sucrose buffer (250 mM Sucrose, 10 mM Tris pH 7.4, 50 µg/ml heparin) as described (Saada *et al*. 2003). For enzymatic activities, mitochondria were pre-incubated with Astaxanthin or DMSO for 5 min on ice prior to assay.

### Blue native electrophoresis for isolation of native organellar complexes

Blue native (BN)-PAGE of mitochondrial fractions was performed according to the method described by (Eubel *et al*. 2003). Mitochondrial pellets were solubilized with n-dodecyl-ß-maltoside (1.5% [w/v]) in ACA buffer (750 mM amino-caproic acid, 0.5 mM EDTA, and 50 mM Tris-HCl, pH 7.0), and then incubated on ice for 30 min. The samples were centrifuged 8 min at 20,000g, and Serva Blue G (0.2% [v/v]) was added to the supernatant. The samples were then loaded onto a native 4% to 16% gradient gel. For non-denaturing-PAGE-Western blotting, the gel was transferred to a PVDF membrane (Bio-Rad) in Cathode buffer (50 mM Tricine and 15 mM Bis-Tris-HCl, pH 7.0) for 16 h at 4°C (const. 40 mA). The membrane was incubated with various antibodies, as indicated for each blot, and detection was carried out by chemiluminescence assay after incubation with an appropriate horseradish peroxidase (HRP)-conjugated secondary antibody.

### SDS-PAGE gel electrophoresis

Organellar protein concentration was determined by the Bradford method (BioRad, Catalog no. 5000201) according to the manufacturer’s protocol, with bovine serum albumin (BSA) used as a calibrator. For immunoassays, an aliquot equivalent to 600 μg *C. elegans* mitochondrial proteins was suspended in sample loading buffer (Laemmli 1970) and subjected to SDS-PAGE (at a constant 100 V). Following electrophoresis, the proteins were transferred to a PVDF membrane (BioRad, Catalog no. 1620177) and incubated overnight at 4^0^C with various primary antibodies (Table S2). Detection was carried out by chemiluminescence assay after incubation with an appropriate horseradish peroxidase (HRP)-conjugated secondary antibody.

### Assessment of mitochondrial respiratory chain enzymatic activities

In-gel activity stains of mitochondrial respiratory complexes I, II and IV was carried out according to (Eubel *et al*. 2005). Following BN-electrophoresis, the gels were washed several times with DDW. For Complex I activity staining, the BN-gels were incubated in CI-activity assay solution (100 mM Tris-HCl, pH 7.4, 0.14 mM NADH, 1 mg/ml NBT solution), for 10 to 30 min until the purple staining of CI bands became visible in the gel. Complex II activity staining was carried out with 50 mM KH_2_PO_4_ buffer (pH 7.4), 84 mM succinate, 0.2 mM PMS and 2 mg/ml NBT solution. For complex IV activity staining, the BN-gels were incubated in 10 mM KH_2_PO_4_ buffer (pH 7.4), 1 mg/ml DAB solution and 0.2 mg cytochrome c. Reactions were stopped in fixing solution containing 15% (v/v) ethanol and 10% (v/v) acetic acid. All steps were carried out at room temperature. Respiratory chain activities and citrate synthase activity assayed by spectrophotometric methods at 37°C(Kirby *et al*. 2007, Shufaro *et al*. 2012). Briefly, complex I was measured as rotenone sensitive NADH-CoQ reductase at 340 nm in the presence of coenzyme Q_1_. Complex II was measured as succinate dehydrogenase (SDH) based on the succinate-mediated phenazine methosulfate reduction of DCPIP (dichloroindophenol indophenol) at 600 nm. Complex II + III was measured as thenoyltrifluoracetone sensitive succinate cytochrome c reductase at 550 nm. Complex III was measured as antimycin sensitive ubiquinol cytochrome c reductase at 550 nm Complex IV. (cytochrome c oxidase) was measured by following the oxidation of reduced cytochrome c at 550 nm. Citrate synthase (CS), an ubiquitous Krebs cycle enzyme, serving as a control, was measured in the presence of acetyl-CoA and oxaloacetate by monitoring the liberation of CoASH coupled to 5’,5-dithiobis (2-nitrobenzoic) acid (i.e., Absorbance at 412 nm).

### Respiration activity

Oxygen consumption (O_2_-uptake) measurements were performed with a Clarke-type oxygen electrode (Oxytherm System, Hansatech Instruments, Norfolk, UK), and the data feed was collected by Oxygraph-Plus software, essentially as described previously (see e.g., (Shevtsov *et al*. 2018). The electrode was calibrated with O_2_-saturated water, and by depletion of the oxygen in the electrode chamber with the addition of excess sodium dithionite. Total respiration was measured at 25°C in the dark, following the addition of 250 µL *C.elegans* suspension from different treatments (± Astaxanthin) and mutant-lines to a 2.25 ml of M9 solution in the respiration chamber. The *C.elegans* suspension was prepared by 1/16 dilution of dense packed worms from 5 plates in M9 buffer. Oxygen consumption was also measured in the presence of various inhibitors, including Rotenone (Rot), Antimycin A (Anti A) and potassium cyanide (KCN).

### Statistical Analyses

Lifespan curves were analyzed by plotting Kaplan-Meier survival curves (Goel *et al*. 2010), and by conducting Log-rank tests (Mantel 1966). Mean lifespan data was compared using Log-rank test with multiple comparisons test, OASIS-2 (Han *et al*. 2016). Differences with P-value smaller than 0.05 were considered as statistically significant.

## Results

### Feeding adult *C. elegans* with Astaxanthin cause significant extension in their lifespan

Synchronized *C. elegans* animals were grown for 5 days at 20°C on NGM plates containing *P. marcusii* cells (expressing Astaxanthin) or in the presence of *P. marcusii* ΔcrtB mutant-line (lacks carotenoids expression). Following 5 days, the intestine of the worms was cleaned by transferring the animals to new NGM plates, containing *E*. *coli* (OP50) cells, for additional 24 hours (Images of *C. elegans* before and after intestine cleaning procedure are provided in Supplementary Figure S2). Under these conditions, the average lifespan of wild-type *C. elegans* (N2), grown on agar plates and fed with *Paracoccus marcusii* (containing on average 716 Astaxanthin per gram of dry material), was about 20% longer (i.e., 24 Days vs. 20 Days) than *C. elegans* fed with *P. marcusii* ΔcrtB mutant that lacks carotenoids (P-value = 0.0038) and about 45% longer (24 Days vs. 17 Days) compared with those of *C. elegans* fed with *E. coli* OP50 cells (P-value = 0.00003) (Fig. 1). Likewise, feeding *C. elegans* with a recombinant *E. coli* cell-line which produces Astaxanthin (pASTA, 146 Astaxanthin per gram of dry material), was also sufficient to significantly increase the lifespan of the worms by about 20% (P-value = 0.0114, Fig. 1B). Feeding *C. elegans* with Astaxanthin extracted from the algae *Haemayococcus pluvialis* and placed on *E. coli* OP50 strain increased the average lifespan by ∼20% (P-value = 0.0173, Fig. 1C). No effect on the lifespan was observed in *C. elegans* fed with *P. marcusii* strains expressing the carotenoids Zeaxanthin (P-value = 0; *P. marcusii* ΔcrtW, Fig. S2 and Table S1), while Lycopene (*P. marcusii* ΔcrtY, Table S1) was found to be toxic to the animals (animals arrested at the L2 stage).

**Figure 1.**
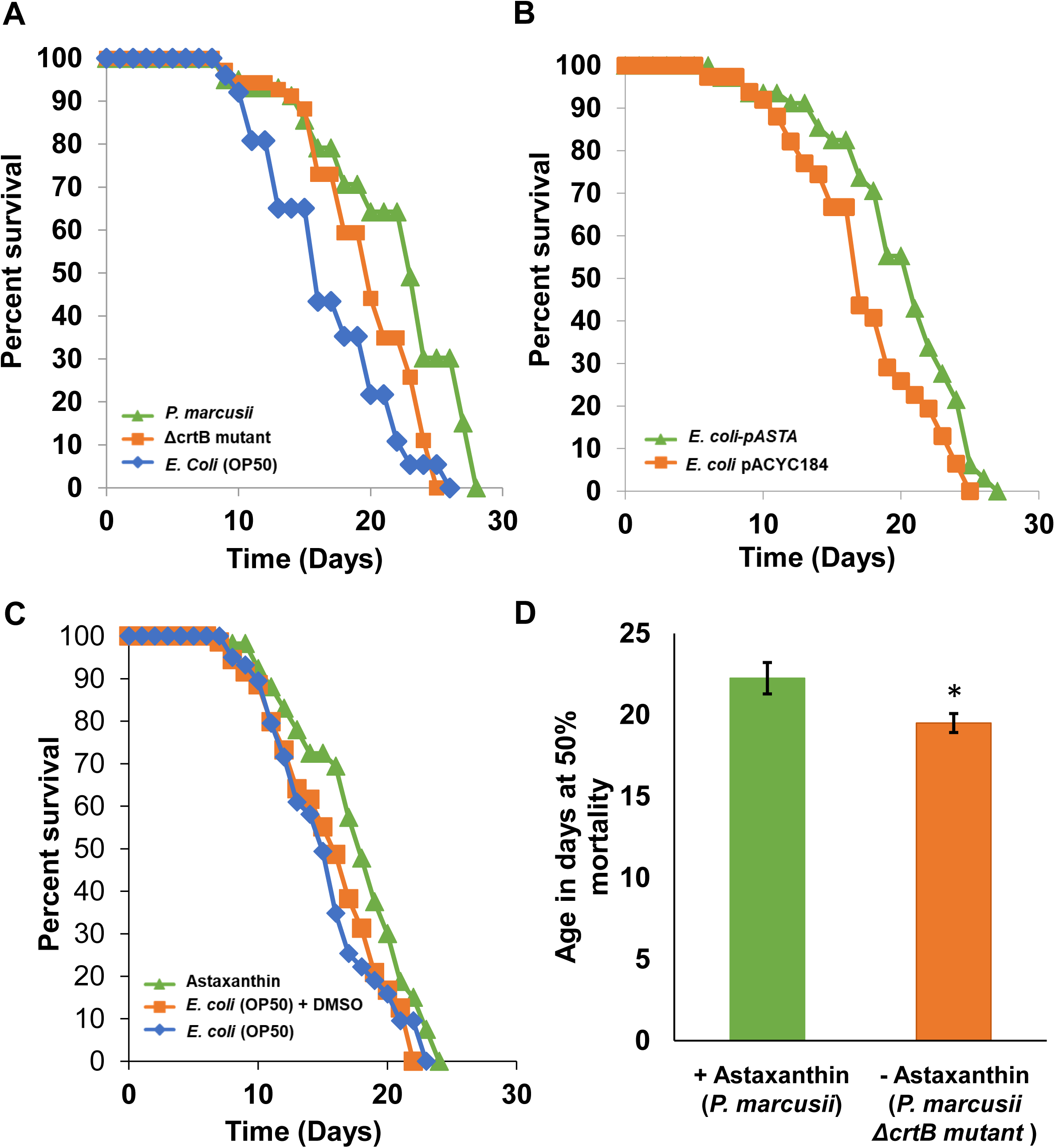
The effects of Astaxanthin on the lifespan of *C. elegans*. Survival curves of *C. elegans* fed with Astaxanthin. Between 70-80 animals were used in each experiment. (**A).** Animals were fed with wild-type *Paracoccus marcusii* (produce Astaxanthin, green), *P. marcusii* ΔcrtB mutant (lacks carotenoids, orange) or *E. coli* (OP50) cells (blue). The results are the average of 4 independent experiments. **(B).** *C. elegans* fed with *E. coli*-pASTA (expressing Astaxanthin, green) and *E. coli* cells containing an empty vector (*E.coli* pACYC184, orange). The mean lifespan (in days) was 19.97 ± 0.74, compared to 17.34 ± 0.69 in the control cells. The results are the average of 5 independent experiments. **(C)**. Animals fed with purified Astaxanthin solubilized with DMSO (green), *E. coli* (OP50) cells (orange) and *E. coli* (OP50) cells + DMSO (blue). The results are the average of 5 independent experiments. **(D)**. The half-life’s of worms fed with *P. marcusii* (produce Astaxanthin, green) and *P. marcusii* ΔcrtB mutant (lacks carotenoids, orange). The results are the average of 7 independent experiments. The values are means of five biological replicates (error bars indicate one standard deviation). Asterisk in panel D indicates a significant difference from control (Student’s T-test, P 0.05). Lifespan curves were analyzed by plotting Kaplan-Meier survival curves (Goel*, et al.* 2010). Mean lifespan data was compared using Log-rank test (Mantel 1966) with appropriate correction for multiple comparisons OASIS-2.

Next, we determined the developmental period where Astaxanthin has the most notable effect on the lifespan of *C. elegans* (see Fig. S3). For this purpose, we used four different feeding conditions. These include animals that were fed from the larval L1 until adulthood with Astaxanthin (i.e., in the presence *P. marcusii*), and later transformed to new plates lacking Astaxanthin (i.e., plates containing *P. marcusii* ΔcrtB) (Fig. S3, L1-Mature +Asta, Adult -Asta). The second condition involved animals from the larva L1 stage until adulthood that were fed with *P. marcusii* ΔcrtB (lacks Astaxanthin) and later transferred to new plates contacting wild-type *P. marcusii* (produces Astaxanthin) (Fig. S3, L1-Mature -Asta, Adult +Asta). In addition, as controls we also used worms that were fed during their entire lifespan on either *P. marcusii* wild-type (Fig. S3, +Asta) or *P. marcusii* ΔcrtB (Fig. S3, -Asta).

As indicated Figure S3, feeding animals that are only at their adult stage (Fig. S3, L1-Mature -Asta, Adult +Asta) with Astaxanthin resulted with an increase in lifespan which was similar to that of animals fed with Astaxanthin throughout their entire life (P-values <0.05). However, no significant change in longevity was apparent when the animals were fed with Astaxanthin from the larval L1 stage until young adulthood, but were avoided from being exposed to the pigment, during their adult stage (Fig. S3, L1-Mature +Asta, Adult -Asta). Similarly, no increase in lifespan was seen when the animals are grown on *P. marcusii* ΔcrtB cells lacking any carotenoids.

We also noted that *C. elegans* animals fed with *P. marcusii* showed a strong red-orange color of the gut (Fig. S4, left panel). Feeding *C. elegans* with *P. marcusii* cells expressing Astaxanthin followed by 24 hours feeding with *E. coli* (OP50) resulted with a strong decolorization of the guts, but some residual red color was still observed in the bod cells (Fig. S4, middle panel). We therefore hypothesize that feeding *C. elegans* at the adult stage with Astaxanthin resulted with the accumulation of Astaxanthin in the cells, an effect that likely correlates with the observed extension in the lifespan.

### The effects of Astaxanthin on *C. elegans* longevity are tightly associated to the mitochondria-aging pathway

There are several known pathways that affect aging in *C. elegans* (i.e., the insulin pathway, the dietary restriction system, and the mitochondrial respiration system) (Amrit*, et al.* 2014, Collins*, et al.* 2008). Here, we performed genetic analyses to determine whether and which of the three known aging pathways might be associated with the prolongevity effect of Astaxanthin. We reasoned that animals with mutations in pathways affected by Astaxanthin would show reduced effects in the presence of the carotenoid on the lifespan extension. Among of the longevity-associated pathway in *C. elegans* is the DAF-16/DAF-2 insulin pathway (Lin *et al*. 2001). We noticed that similarly to the-wild-type animals, Astaxanthin caused an (additional) increase in the lifespan of the *daf-2 (e1370)* (P-value = 0.001) and *daf-16 (mu86)* (P-value = 0.000046) mutant animals (Fig. 2). Similarly, feeding *C. elegans glp-1 (or178)* mutants (Libina *et al*. 2003, TeKippe and Aballay 2010) with Astaxanthin resulted in additional extension of the lifespan (P-value = 0.000024, Fig. 2). We therefore concluded that Astaxanthin effect on *C. elegans* lifespan are independent of the insulin/IGF signaling pathway. Another well-characterized pathway of lifespan extension involves dietary restriction (Lakowski and Hekimi 1998). Here too, feeding *eat-2 (ad1116)* mutant animals with Astaxanthin resulted with an extended lifespan (P-value = 0.011, Fig. 2), indicating that Astaxanthin effect on *C. elegans* lifespan are also independent of the dietary restriction mechanism.

**Figure 2.**
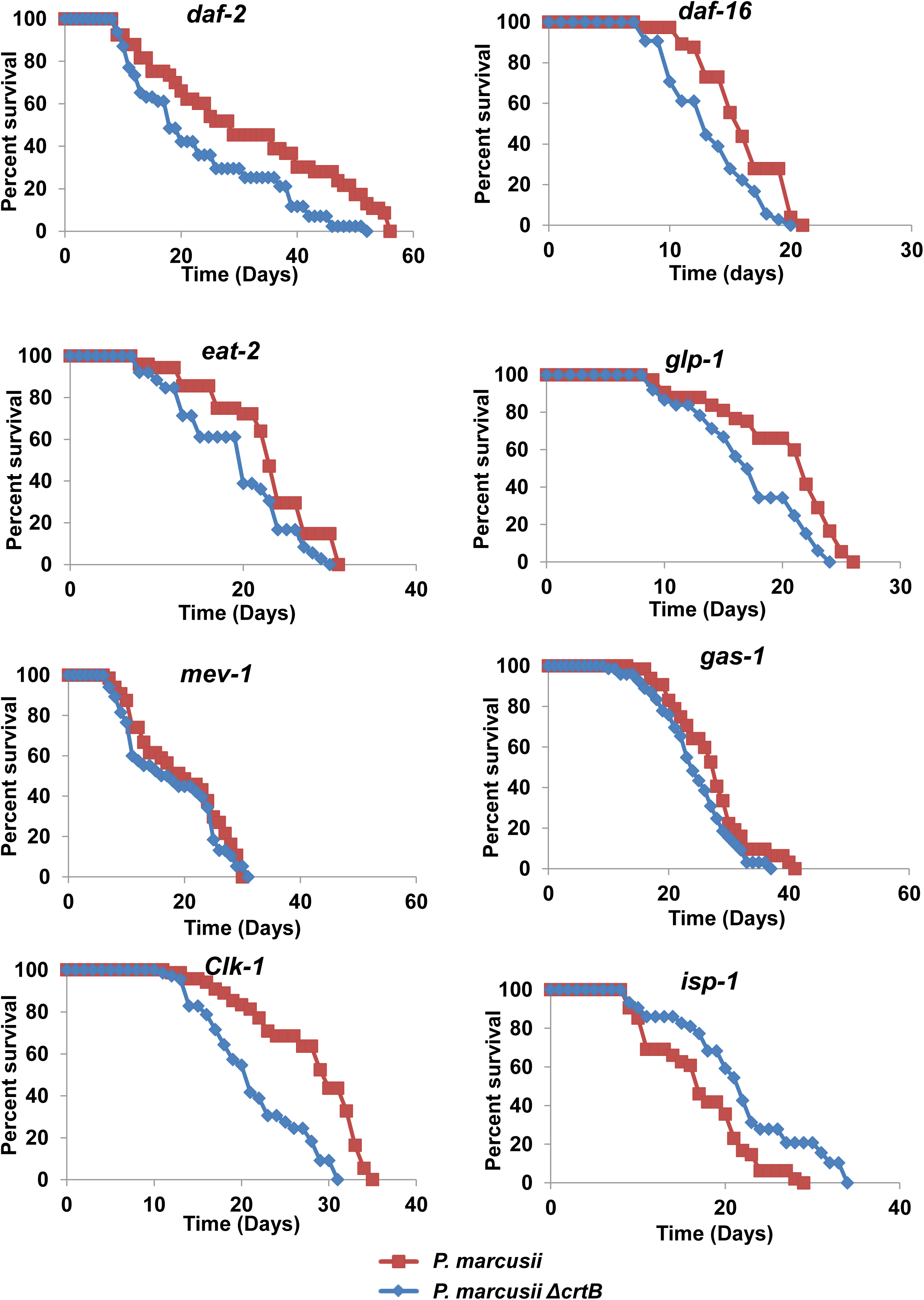
The effects of Astaxanthin on the longevity of *C.elegans* mutated in different life-extending genes. Survival curves of nematodes mutated in genes that regulate lifespan through different pathways supplemented with Astaxanthin. Worms mutated in the *daf-2(e1370)*, *daf-16(mu86)*, *eat-2(ad1116)*, *glp-1(or178)*, *isp-1(qm150)*, *mev-1(kn1*), *gas-1(fc21)* or *clk-1(e2519)* were fed with *P. marcusii* (produce Astaxanthin, red), *P. marcusii* ΔcrtB mutant (lacks carotenoids, blue). Nematode survival was calculated with the Kaplan-Meier method (Goel*, et al.* 2010), and differences in survival rates were tested for significance using the log-rank test (Mantel 1966), with appropriate correction for multiple comparisons OASIS-2.

In addition to the insulin/IGF signaling and dietary restriction pathways, life-extending in animals also involves the mitochondrial respiration system (i.e., mitochondrial dysfunctions resulting with extended lifespan) (Murakami and Murakami 2005). It is postulated that mutations in genes encoding components of the mitochondrial electron transport chain (ETC) lower the levels of ROS and thereby may increase the lifespan of the animals (Murakami and Murakami 2005). Electron leakage from complex III is suggested to be a major contributor to the generation of ROS within the mitochondria (Brand 2010, Chen et al. 2003, Quinlan et al. 2013). To test whether reduced respiration activities may influence the prolongevity-mediated effects by Astaxanthin, we analyzed the lifespan of *C. elegans* mutants in genes encoding respiratory-related proteins. Mutants in the Rieske iron-sulfur protein (RISK) of complex III (i.e., *isp-1* mutants, also denoted as *qm150)*, showed reduced (shortened) lifespan when they were treated (fed) with Astaxanthin (P-value = 0.000011, Fig. 2). These results strongly suggest that the Astaxanthin-mediated longevity effects involve complex III. The effects of Astaxanthin on the lifespan was further analyzed in *C. elegans* mutants affected in other mitochondrial proteins. *CLK-1* encodes a demethoxyubiquinone hydroxylase enzyme required for ubiquinone biosynthesis. *C. elegans clk-1* mutants show a pleiotropic phenotype that includes slowed development and aging (Branicky *et al*. 2006). However, the data showed that Astaxanthin-mediated longevity was not affected in the *clk-1* (e2519) animals (P-value = 1.1 x 10^-7^, Fig. 2). Feeding *C. elegans gas-*1 (*fc21*) (affected in complex I) or *mev-*1 (*kn1*) (affected in complex II) mutants with Astaxanthin, resulted with mild extensions in longevity (P-values 0.0024 and 0.2831, respectively). Taken together, these data indicate that the Astaxanthin-extended lifespan effects we see in *C. elegans* rely on the level or activity of respiratory complex III, and maybe to some extend on other respiratory-mediated functions.

### Astaxanthin affects the biogenesis of the native respiratory complex III, *in vitro*

The Astaxanthin-mediated effects we see suggest that Astaxanthin increases the lifespan of *C. elegans* by influencing the biogenesis or function of the respiratory system via CIII and maybe CI and CII as well (Fig. 2). To analyze whether Astaxanthin directly affect the biogenesis or activities of these respiratory complexes, we used *in vitro* analyses with crude mitochondria preparations (i.e., ‘*in-organello*’ assays), obtained from both animals and plants. These include *C. elegans*, rodents, human and cauliflower mitochondria (*Brassica oleracea* var. botrytis) (see Materials and Methods). For this purpose, Astaxanthin was extracted from the algae *Haematococcus pluvialis*, solubilized in 5% DMSO and then incubated with the mitochondria for 30 min on ice. The integrity of native respiratory complexes in *C. elegans*, cauliflower, rodents and human, was investigated by BN-PAGE analyses (Fig. 3 and Fig. S5).

**Figure 3.**
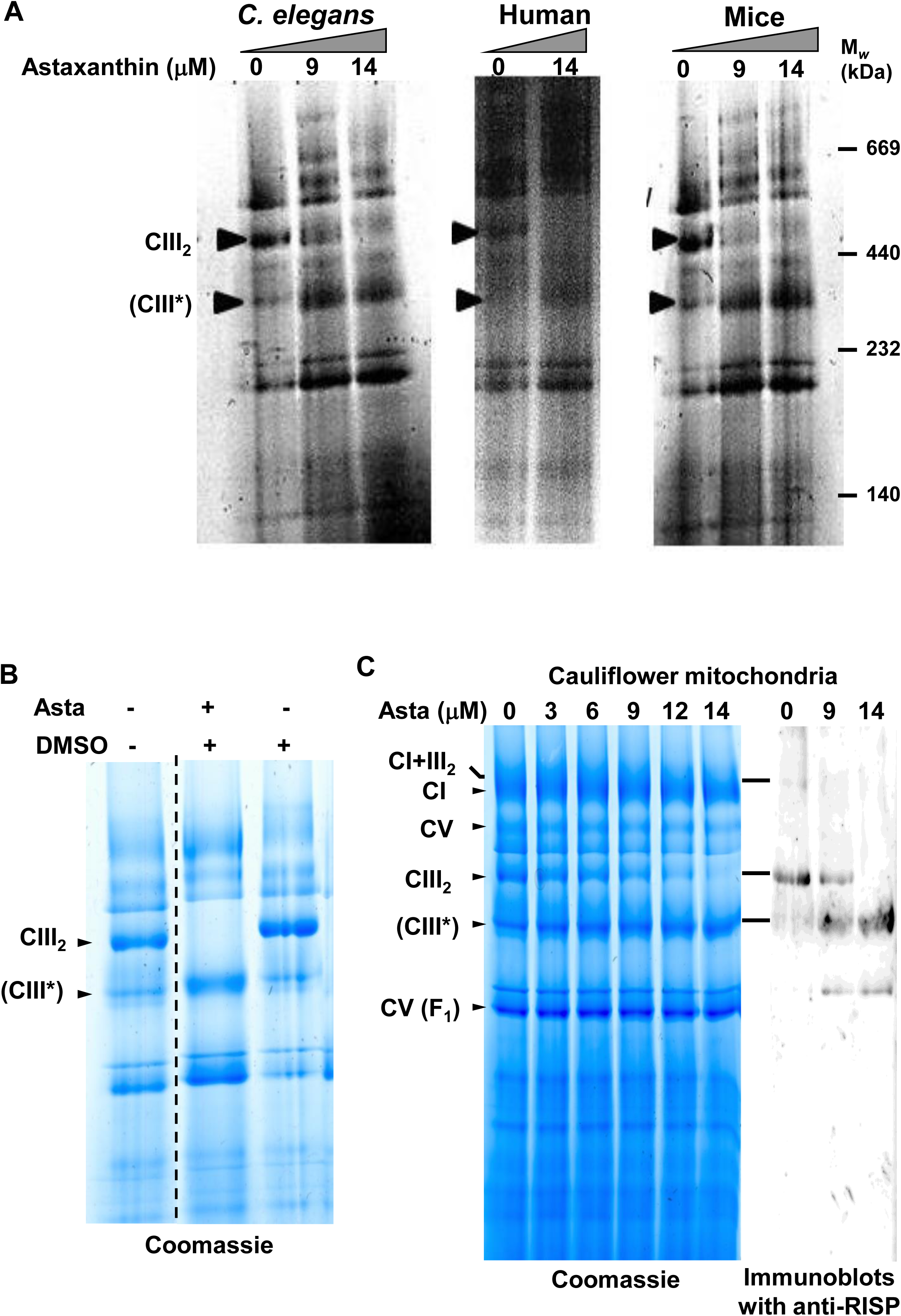
The Effect of Astaxanthin on isolated plant mitochondria. **(A)**. Astaxanthin was added at different concentrations (i.e., 0, 9 and 14 μM) to crude mitochondria preparations, obtained from *C. elegans*, human and mice. Native organellar complexes were separated by BN-PAGE, followed by Coomassie Blue staining. The position of complex III, which partially disassembled in the presence of Astaxanthin, is indicated by arrows. **(B).** The effects of Astaxanthin on various respiratory complexes in cauliflower mitochondria was analyzed by BN-PAGE assays. **(C)** Increasing amounts of Astaxanthin (as indicated in the gel) were added to isolated mitochondria of cauliflower. Arrows indicate to native complexes I, III and V. The effects of Astaxanthin on CIII was analyzed by western-blot analysis with antibodies raised against plant-specific RISP (Rieske iron sulfur) protein. The blots indicate that Astaxanthin cause a reduction in the levels of the holo-complex III and supercomplex I+III, and to the appearance of protein-bands of lower molecular-masses.

Figure 3A shows the effects of increased Astaxanthin concentrations (i.e., 0, 9 and 14 μM) on native respiratory complexes in animal mitochondria. A decreased level of a high-molecular mass band (about 500 kDa), which we expected to corresponds to a native complex III dimer (i.e., holo-CIII_2_), was observed in *C. elegans* mitochondria treated with Astaxanthin. Notably, this Astaxanthin-mediated reduction in the ∼500 kDa band, was followed by the appearance of a ∼350 kDa band, which may corresponds to a partially disassembled CIII particle (i.e., CIII*). BN-PAGEs of crude mitochondria preparations obtained from rodents and human treated Astaxanthin showed the same patterns (i.e., reduced intensities of the ∼500 kDa complex and an accumulation of ∼350 kDa particles) (Fig. 3A). To analyze whether similar effects may occurs in vivo, we analyzed the integrity of respiratory complexes in *C. elegans* fed with Astaxanthin. For this purpose, young adult worms were grown for 7 days in the presence of wild-type *P. marcusii*, washed several times to remove bacterial contaminants and crude mitochondria preparations were obtained, essentially as described previously (Grad*, et al.* 2007). Organellar proteins from *E.coli* (-Asta) and *P. marcusii* (+Asta) fed *C. elegans* were separated by under native conditions. The BN-PAGE assays indicated that the absorption or uptake of Astaxanthin (Fig. S4) also resulted with a reduction in the 500 kDa band and the accumulation of the 350 kDa particles (Fig. S5).

We next analyze the effects of Astaxanthin on mitochondria isolated from cauliflower inflorescences, as this tissue allows the purification of large quantities of highly-pure mitochondria preparations (Keren*, et al.* 2009, Neuwirt*, et al.* 2005, Sultan*, et al.* 2016). Mitochondria respiratory complexes were then separated by BN-PAGE analysis (Fig. 3B and Supplementary Fig. S6). Arrows indicate to native respiratory complexes CI (∼1,000 kDa), a CIII dimer, (CIII_2,_ about 500 kDa), CIV (∼220 kDa) and CV (∼600 kDa) in cauliflower mitochondria. In plant’s mitochondria as well, the addition of Astaxanthin led to a notable reduction in the levels of the 500 kDa corresponding to CIII_2_, followed by the accumulation of a lower (about 300 kDa band) corresponding to sub-CIII particle (CIII*) (Fig. 3B). These were correlated with the levels of Astaxanthin added to the isolated mitochondria (Fig. 3C). BN-PAGEs followed by immunoblot analyses with antibodies raised to RISP subunit (iron-sulfur protein of complex III), indicated that both the ∼350 kDa and 500 kDa particles contain RISP (Fig. 3 and Fig. S6), strongly supporting that these particles correspond to holo-CIII2 and sub-CIII*. The Astaxanthin-mediated effects (i.e., decreased CIII_2_ levels and increased amounts of the 350 kDa band) seem specific to Astaxanthin, as mitochondria treated with DMSO alone had no obvious effects on the levels or integrity of respiratory complexes I, III and V in cauliflower mitochondria (Fig. 3B), while the addition of β-Carotene or Zeaxanthin had no obvious effects on the integrity of the organellar complexes (Fig. S5B). Mass-spectrometry (LC-MS/MS) analyses of gel slices corresponding to holo-CIII_2_ (upper band) and CIII* particles (lower band) obtained from human (Table S3) and plant (Table S4) mitochondria, indicated the presence of numerous subunits of CIII (see Table 2). The LC-MS/MS data also indicated the presence of many other peptides that are corresponding to CI (mainly), CIII, CIV, CV and various other known mitochondrial proteins. These may be associated with the CIII particles. However, it is anticipated that these proteins rather correspond to various sub-particles of respiratory complexes that are migrating with similar masses to those of the holo-CIII_2_ and CIII*, as only the immunoblots with RISP analyses indicated a correlation with the reduction in the CIII dimmer and to increased levels of the lower (350 kDa) band (Figs. 3C and S6). As indicated by immunoblots, the presence of RISP protein in the holo-CIII_2_ and CIII* of both human and cauliflower mitochondria, was also supported by the LC-MS/MS data (Table 2 and Supplementary Tables S3 and S4).

**Table 1.**
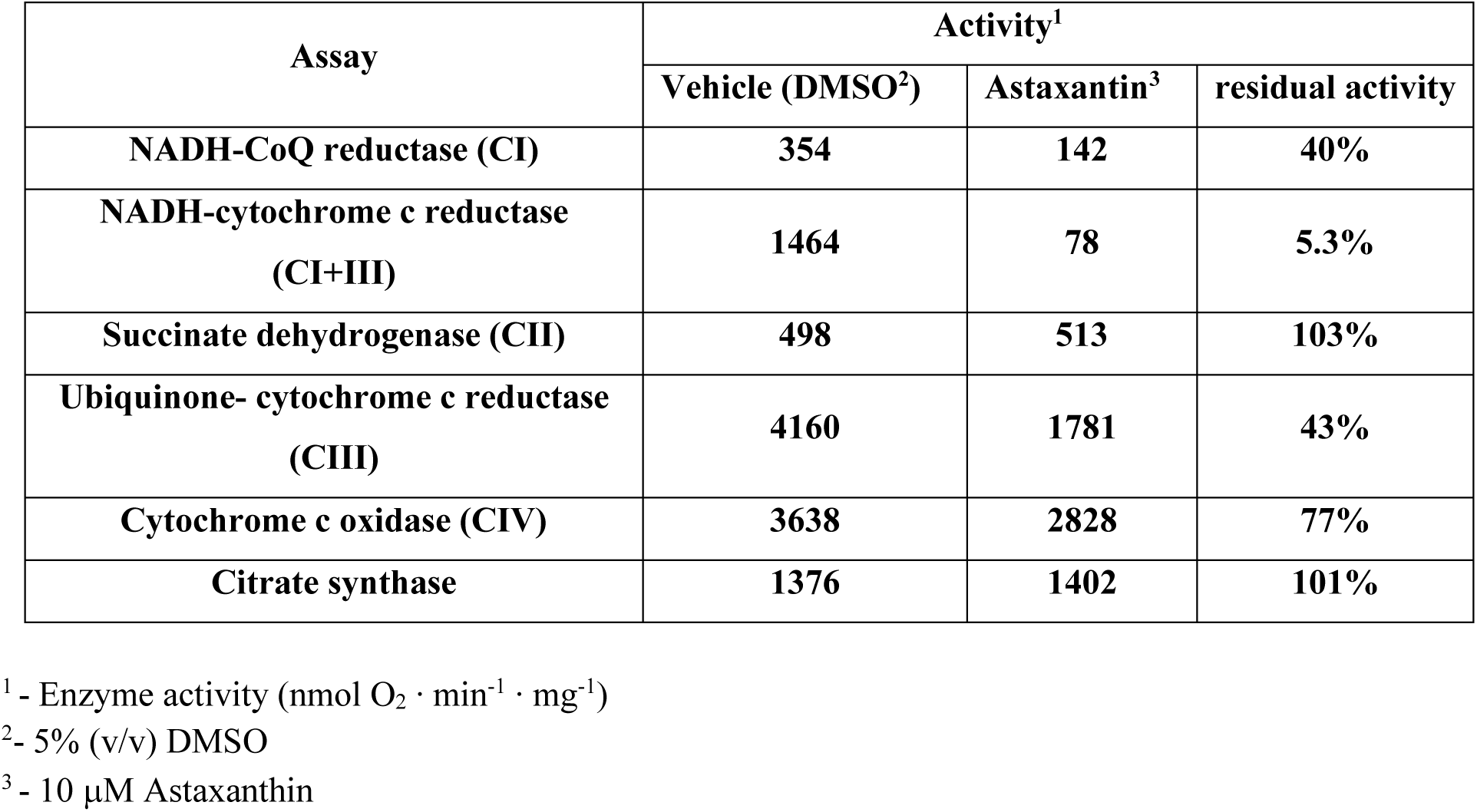
The effects of Astaxanthin on the enzymatic activities of mouse heart mitochondrial respiratory complexes.

| Assay | Activity<sup>1</sup> |  |  |
| --- | --- | --- | --- |
|  | Vehicle (DMSO<sup>2</sup>) | Astaxanthin<sup>3</sup> | residual activity |
| NADH-CoQ reductase (CI) | 354 | 142 | 40% |
| NADH-cytochrome c reductase (CI+III) | 1464 | 78 | 5.3% |
| Succinate dehydrogenase (CII) | 498 | 513 | 103% |
| Ubiquinone- cytochrome c reductase (CIII) | 4160 | 1781 | 43% |
| Cytochrome c oxidase (CIV) | 3638 | 2828 | 77% |
| Citrate synthase | 1376 | 1402 | 101% |

<sup>1</sup> - Enzyme activity (nmol O<sub>2</sub> · min<sup>-1</sup> · mg<sup>-1</sup>)
<sup>2</sup>- 5% (v/v) DMSO
<sup>3</sup> - 10 µM Astaxanthin

**Table 2.** Summary of mass-spectrometry (MS) of the ∼500 kDa (higher) and ∼350 kDa (lower) particles in animals (human) and plant (cauliflower) mitochondria, treated with Astaxanthin.

|  | Human |  | Cauliflower |  |
| --- | --- | --- | --- | --- |
| Associated respiratory complex | No. of subunits identified in the 500 kDa band | No. of subunits identified in the 350 kDa band | No. of subunits identified in the 500 kDa band | No. of subunits identified in the 350 kDa band |
| CI (NADH-dehydrogenase) | 40 | 20 | 11 | 9 |
| CII (succinate dehydrogenase) | 2 | 2 | 2 | 2 |
| CIII (Cytochrome c reductase) | 13 | 12 | 11 | 10 |
| Rieske iron-sulfur protein | + | + | + | + |
| CIV (Cytochrome c oxidase) | 15 | 11 | 1 | 1 |
| CV (ATP-synthase) | 13 | 11 | 8 | 7 |

In addition to CIII, Astaxanthin may also affects the accumulation of CI (using anti-CA2 antibodies) and the ATP-synthase (CV, anti-ATPA antibodies) in cauliflower mitochondria (Fig. 3C and Fig. S6). BN-PAGE analyses followed by immunoblots corresponding to CI (with anti-CA2 antibodies) and ATP-synthase (CV, anti-ATPA antibodies) indicated to reduced holo-CI (∼1,000 kDa) and a ∼1,500 kDa particle, corresponding to a ‘supercomplex’ containing CI and CIII_2_ (i.e., CI+CIII_2_), which was also apparent in the immunoblots with anti-RISP antibodies (Figs. 3C and S6). Yet, the pigment has no obvious effect on the levels and biogenesis of the native complex V (Fig. 3C and Fig. S6).

The steady-state levels of various mitochondria proteins in Astaxanthin-fed *C. elegans* was assayed by immunoblot analyses, with antibodies generated to various respiratory complexes, including CI (C25H3.9), CIII (ISP-1), CIV (CTC-1) and CV (ATP-5) and cytochrome C (CYC-1) (see Table S2). In each assay, the steady-state levels of different proteins in the Astaxanthin-fed animals were compared to those of untreated wild-type animals of the same age at different total protein loadings (i.e., 0.15 up to 1.5 times) (Fig. 4). The levels of various mitochondrial proteins, including C25H3.9 (CI), CTC-1 (CIV) and ATP-5 (CV) did not changed significantly between the control and Astaxanthin-fed animals or the *C. elegans isp-1* mutants (Fig. 4B). A reduction in the signal of cytochrome C (CYC-1) was observed in *C. elegans isp-1* mutants, but not in animals fed with Astaxanthin. We also noticed a notable reduction in RISP signals in both *isp-1* animals and wild-type *C. elegans* fed with Astaxanthin. While a reduction in RISP is expected in the case of the *isp-1* mutant-line (Fig. 4, ISP-1 panel), it remains unclear why a notable reduction in RISP is seen in animals fed with Astaxanthin, whereas the protein remains intact in the ‘*in organello*’ assays. A reasonable explanation is that when the worms are fed with Astaxanthin they undergo continues disassembly of CIII (or the suppercomplex of CI-CIII). Under these conditions, various complex III (and possibly CI) subunits, including the RISP protein, may become more susceptible to proteolysis by organellar proteases and peptidases, while much lower degradation rates are expected under the in vitro conditions (i.e., short incubations on ice).

**Figure 4.**
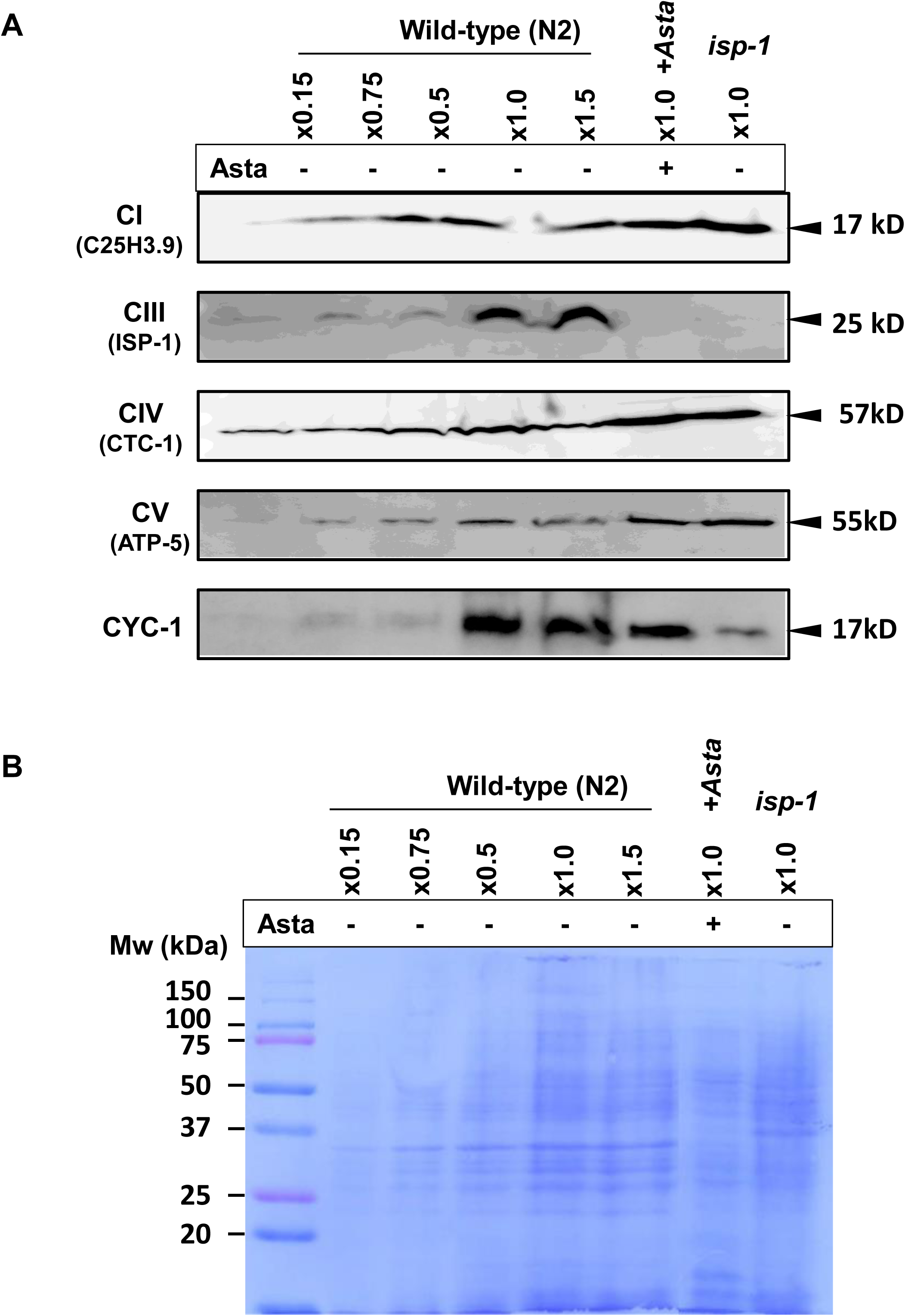
Changes in the mitochondria RISC complex of *C. elegans* grown on bacteria expressing Astaxanthin. (**A).** Western-blot analysis of crude mitochondria obtained from wild-type *C. elegans* at different concentrations*, C. elegans* fed with Astaxanthin (+Asta), and *C. elegans isp-1* mutant-line. Antibodies against NUFB5 (a subunit of the mitochondrial NADH dehydrogenase (ubiquinol) of the mitochondrial complex I (ortholog in *C. elegans* is termed as C25H3.9), Rieske iron sulphur protein (ISP-1) subunit of the mitochondrial complex III, COX1 (cytochrome c oxidase subunit 1) of the mitochondrial complex IV (the ortholog in *C. elegans* is termed as CTC-1), ATP5A (ATP synthase subunit alpha subunit) of mitochondrial complex V (the orthologue in *C. elegans* is ATP-5), and Cyt C (cytochrome c reductase; the ortholog in *C. elegans* is CYC-1), were used in immunoblotting assays. (**B).** SDS-PAGE followed by Coomassie staining of isolated mitochondria from *C. elegans* fed with Astaxanthin (+Asta). Controls included crude mitochondria isolated from wild-type *C. elegans* that were not fed with Astaxanthin, and mitochondria isolated from the *C. elegans isp-1* mutant-line.

### *C. elegans* treated with Astaxanthin display altered respiratory functions

Analyses of native mitochondrial complexes, using denaturing PAGE profiles, suggested to alter CIII and maybe CI biogenesis in *C. elegans* fed with Astaxanthin. To determine whether the respiratory activity was affected in animals fed with Astaxanthin, we monitored the oxygen (O_2_) uptake rates of non-treated versus Astaxanthin-fed wild-type *C. elegans* and *isp-1* mutants, using a Clark-type electrode (Hansatech Instruments, Norfolk, UK). For this purpose, synchronized late L4 worms were collected and washed several times with M9 buffer. Oxygen consumption was monitored in the worms in a sealed chamber (see e.g., (Cohen *et al*. 2014, Keren*, et al.* 2012, Shevtsov*, et al.* 2018, Zmudjak *et al*. 2013). Each respiration chamber contained 250 µl of packed *C. elegans* suspension diluted 10 times in M9 buffer. These measurements indicated that *C. elegans* fed on Astaxanthin and the *isp-1* mutants have reduced respiration activities (i.e., 15.03 ± 3.03 and 14.33 ± 2.88 nmol O_2_·min^-1^·mg^-1^ total protein), respectively, compare to those of the non-treated wild-type animals (i.e., 28.83 ± 3.07 nmol O_2_·min^-1^·mg^-1^ total protein) (Fig. 4).

Here, we also measured the O_2_-uptake rates in the presence of various inhibitors, including Rotenone (Rot), which inhibits complex I electron transport, the complex III inhibitor Antimycin A (Anti A) and Potassium cyanide (KCN) that blocks electron transport from complex IV to oxygen. As shown in Figure 4, both Rot and Anti-A have noticeable effects on the respiratory activities of the non-treated wild-type, compared to those of *isp-1* mutant or *C. elegans* fed with Astaxanthin (55% ± 2.95 and 66% ± 0.70 compared to 34% ± 0.99 O_2_·min^-1^·mg^-1^ total protein, respectively). In contrast, KCN had similar effects on the O_2_-uptke rates of wild-type worms, worms fed on Astaxanthin and the *isp-1* mutants (85% ± 5.18 and 90% ± 2.35, and 95% ± 0.19, O_2_·min^-1^·mg^-1^ total protein, respectively) (Fig. 4). These results further indicate to altered respiratory CI- and CIII-mediated activities.

### Astaxanthin inhibits enzymatic activities of mammalian complexes I and III

The BN-PAGE (Fig. 3 and Suppl. Figs. S4-S6), LC-MS/MS (Table S4) and the respiration activity data (Fig. 4) indicate that Astaxanthin has a strong effect on the biogenesis and functions of CIII, and presumably affects CI and the supercomplex I+III, as well. These data were further supported by analyses of the enzymatic activities of native respiratory complexes in the presence or absence of Astaxanthin. Table 1 shows the effects on the respiratory functions of mitochondrial complexes isolated from mouse heart that were pre-incubated with DMSO alone (Mock) or with purified Astaxanthin dissolved in DMSO (see Materials and Methods). The biochemical assays indicated to significant decreases in the activities of both complexes I and III. The inhibitory effect of Astaxanthin was even more pronounced in enzymatic activity assays of CI+CIII, which approached the respiratory assay sensitivity limits. These analyses also indicated that complexes II, IV and citrates synthase enzyme where not, or only slightly, affected by the addition of Astaxanthin to animals mitochondria.

## Discussion

The nematode *Caenorhabditis elegans* is a well-established animal system to study organismal aging, and for the characterizing and identifying new drugs that can extend lifespan and improve the quality of old age in animals. Here, we used *C. elegans* to study the molecular basis of the anti-aging effects of Astaxanthin consumed by various organisms. Our results show that Astaxanthin extends *C. elegans* lifespan by 20%-40%. The effects on the life span seem specific to Astaxanthin, as other carotenoids tested here, including Zeaxanthin and Lycopene had no lifespan extension effects. In addition, the anti-aging role of Astaxanthin had no obvious effects on the reproduction, age pigmentation and locomotion of the animals. The life-extension phenotype, induced by Astaxanthin, relies mainly on *isp-1* mutants that are affected in mitochondria respiratory complex III, but not on *gas-1* (complex I) and *clk-1* (ubiquinol biosynthesis), nor on other aging-related pathways (i.e., the insulin/IGF-1 and dietary restriction pathways), as indicated by lifespan measurements of *daf-2*/*daf-16*, *eat-2* or *glp-1* mutants fed with Astaxanthin.

Remarkably, Astaxanthin added to isolated mitochondria causes a reduction in the levels of holo-complex-III_2_ and suppercomplex I+III, and to the appearance of lower molecular-mass particles that correspond to sub-CIII particles. Accordingly, BN-PAGE and LC-MS/MS analyses indicated the presence of various CIII subunits in both the high and lower mass particles. The presence of various complex I subunits in these gel-bands, may relate to the strong association of complexes I and III, which together with complex IV also form a stable supercomplex known as the Respirasome (I_1_III_2_IV_1_) (Schafer *et al*. 2006). Alternatively, it remains possible that sub-complex I, IV and V (natively formed or disassemble during mitochondria preparations or gel-run) may migrate with similar masses under the experimental conditions. Together, these analyses indicate that Astaxanthin affects the assembly or biogenesis of complex III and the CI-CIII_2_ suppercomplex, whose functional roles and assembly are under investigation (Signes and Fernandez-Vizarra 2018). As CIII biogenesis / disassembly are observed in both animal and plant mitochondria (Figs. 3, S5 and S6), the data imply to a universal effect caused by Astaxanthin to CIII, in all eukaryotes.

Reduced accumulation of CIII is expected to affect the respiratory functions of *C. elegans* cells fed with Astaxanthin. The O_2_-uptake rates of worms fed on Astaxanthin or *isp-1* mutants were reduced by about 2x folds (Fig. 5), compared to those measured in the non-treated (control) *C. elegans*. Similarly, the oxygen consumption of Astaxanthin-treated mitochondria, isolated from mouse cells was lowered by more than 50%, and to reduced activities of the respiratory complexes I and III. Currently, it remains unclear how Astaxanthin affects the assembly or biogenesis of complex III in plants and animals. It is possible that Astaxanthin interacts with complex III and thereby interfere with the assembly process or destabilizes the integrity of the native complex. Due to its skeletal properties (long polyene chain as well as two oxygenated β-ionone-type rings), Astaxanthin seems to share some chemical equivalents with ubiquinol (contains a single benzylquinone head-group ring). Astaxanthin may therefore interfere with the association of ubiquinol with CIII, affecting RISP stability, the assembly state of the holo-CIII or the integrity of the CI-CIII_2_ suppercomplex. Another possible effect of Astaxanthin may involve cardiolipin, a unique phospholipid that is an important component of the inner-mitochondrial membrane. Crystal structures show that cardiolipin (CL) is tightly associated with CIII in vivo (Lange *et al*. 2001) and affects the assembly of various respiratory complexes (Paradies *et al*. 2014). The absorbtion of Astaxanthin (in vivo or in vitro) may interfere with the association of CL with CIII and thereby affects its biogenesis and function. However, such speculations require further investigation.

**Figure 5.**
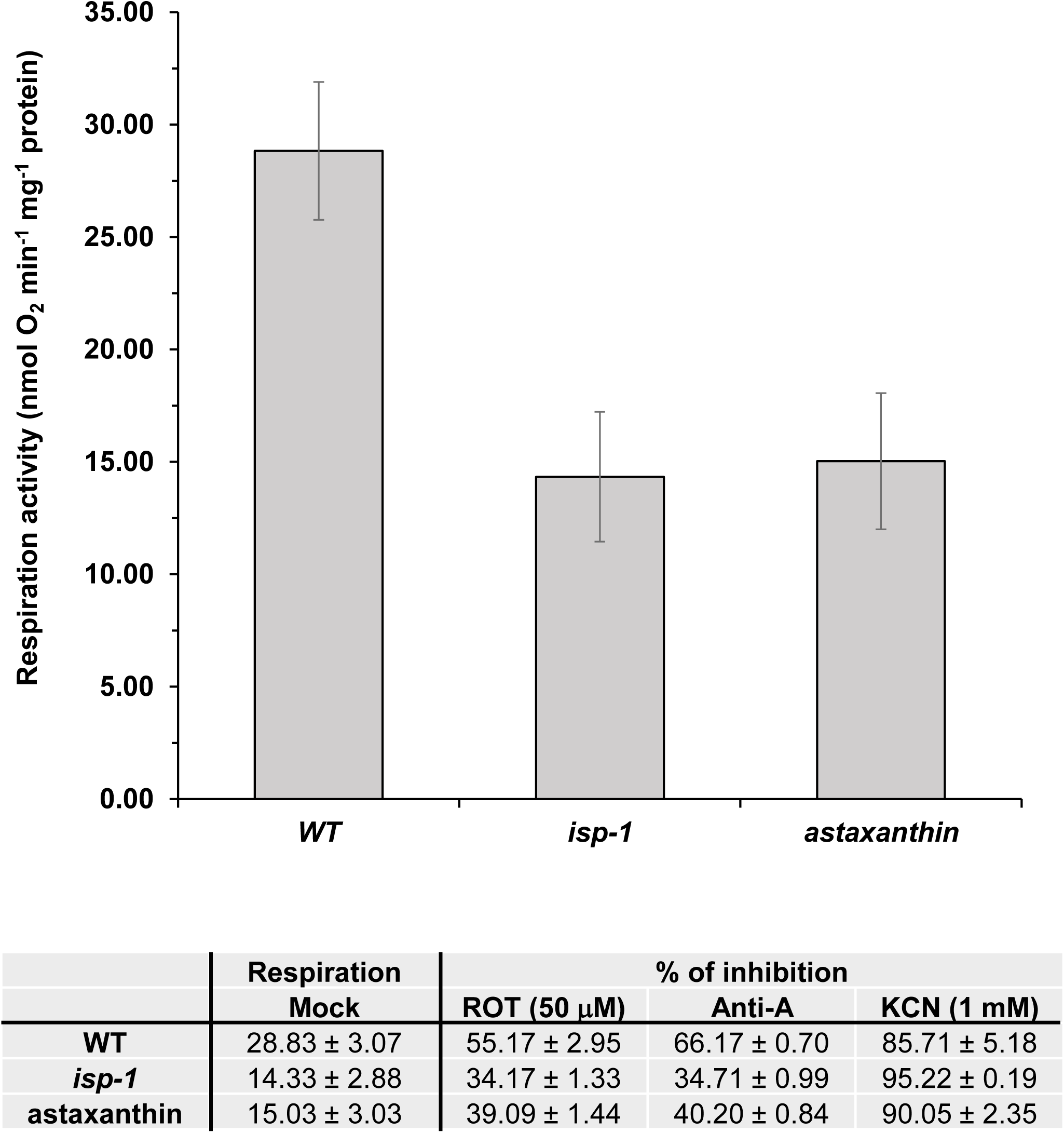
Respiration activities in *C. elegans* wild-type and *isp-1* mutants fed with Astaxanthin. O_2_-uptake rates of wild-type, Astaxanthin-fed animals and *isp-1* mutants were analyzed with a Clark-type electrode as described previously (Cohen*, et al.* 2014). For each assay, equal amounts of dense packed worms from 5 different plates were submerged in 2.25 mL M9 buffer and applied to the electrode in a sealed glass chamber in the dark. O_2_-uptake rates were measured in the absence (Control) or in presence of rotenone (+ROT, 50 μM), Antimycin A (Anti A, 50 μM) and KCN (1 mM) which inhibit complexes I, III and IV activities (respectively). The values are means of four biological replicates. Error bars indicate one standard deviation.

At this point, it is also not clear to us why RISP was reduced in *C. elegans* fed with Astaxanthin, but seem to remained intact in the ‘*in organello*’ assays of mitochondria treated with Astaxanthin (Figs. 3 and S4). The data indicate that Astaxanthin affects the assembly of CIII, as evident by the appearance of lower molecular mass particles corresponding to sub-CIII particles in the mitochondria. Disassembly of CIII is expected to affect the stability of various CIII subunits, including RISP, leading to their degradation over time, whereas under *in organello* assays (i.e., short incubations on ice; see Materials and Methods), RISP (or various other CIII subunits) remain intact and are less prone for degradation by the organellar proteases.

Mitochondria are major sites for energy metabolism and ATP production in eukaryotic cells, through the oxidative phosphorylation pathway. In light of the expected significance of mitochondria functions to animal and plant physiology, why does the reduced CIII activity and level in Astaxanthin-fed animals or the *isp-1* mutants results with an extended life span? Currently, we cannot provide a definitive explanation, but we speculate that these phenotypes relate to reduced mtROS production. ROS are known to induce oxidative damages of fatty acyl chains of membrane lipids, by ‘lipid-peroxidation’ (Tsuchiya *et al*. 1994). Altered membrane integrity has many pathological consequences in human, such as neurological diseases, inflammations, diabetes, cancer, and aging. Astaxanthin is a strong antioxidant that was recently suggested to reduce mtROS protection in *C. elegans* and other animals (Kuraji *et al*. 2016, Liu*, et al.* 2016, Wu *et al*. 2015). Our data indicate to an additional mechanism by which Astaxanthin affects the life span of animals. The association of Astaxanthin with CIII destabilizes the native complex, resulting with lower molecular mass CIII intermediates and reduced respiration. Lowered respiration activities are likely associated with reduced rates of ‘toxic’ electron leakage from complex III, a major contributor to the generation of mtROS (Brand 2010, Chen *et al*. 2003, Quinlan *et al*. 2013), and thereby can protect the cells (and organelles) from oxidative damages. Accordingly, the aging related mechanism through mitochondria activities have been previously related to ROS production (Murakami and Murakami 2005, Quinlan*, et al.* 2013).

In summary, our data provide with novel insights into the molecular mechanism of longevity extension mediated by Astaxanthin in *C. elegans*. Biochemical and genetic studies indicate that Astaxanthin affects complex III biogenesis and activity, and may increases longevity by protecting the animal cells against ROS and oxidative stresses, due to complex III-associated ‘electron leakage’ during respiration. These effects may be universal among different eukaryotes, as similar effects on CIII biogenesis or assembly were noted in plants, warms, rats and human. Our data may also become useful for the development of multi-target drugs designed to CIII in order to prolong the lifespan of animals.

## Acknowledgments

We thank Yuval (Yukon) Stein (The Hebrew University, Jerusalem, Israel) for his assistance with organellar preparations. This work was supported by grants to O.O.B from the ‘Israeli Science Foundation’ (ISF grant no. 741/15).

## Supplementary data

**Table S1.** Carotenoid composition in bacterial strains used in this work.

| **Name** | **Carotenoid composition** |
| --- | --- |
| *Escherichia coli* WT | None |
| *E. coli* pASTA | Astaxanthin |
| *E. coli* pACYC184 | None |
| *Paracoccus marcusii*<br>(*P.marcusii*) wild-type | Largely Astaxanthin<br>Minor: Phoenicoxanthin Adonixanthin,<br>3'-Hydroxyechinenone |
| *P. marcusii* ΔcrtB | None |
| *P. marcusii* ΔcrtW | Zeaxanthin |
| *P. marcusii* ΔcrtY | Lycopene |

**Table S2.** List of antibodies used for the analysis of astaxanthin effects on C. elegans lifespan.

| **Antibody** | **Protein I.D.** | **origin** | **serum** | **dilution** | **Reference / source** |
| --- | --- | --- | --- | --- | --- |
| Atp-A | Mitochondrial ATP-synthase subunit $\alpha$ | *Zea mays* | Mouse (monoclonal) | 1/5,000 | (Michael *et al.* 1993) |
| ATP-5 | ATP synthase subunit, mitochondrial | *Caenorhabditis elegans* | Rabbit (polyclonal) | 1/1,000 | Thermo Fisher Scientific |
| C25H3 | Uncharacterized protein, mitochondrial respiratory complex I | *Caenorhabditis elegans* | Rabbit (polyclonal) | 1/1,000 | Sigma-Aldrich Co. LLC |
| CA2 | γ-carbonic anhydrase-like subunit 2 | *Arabidopsis thaliana* | Rabbit (polyclonal) | 1/1,000 | (Perales *et al.* 2005, Sunderhaus *et al.* 2006) |
| CTC-1 | Cytochrome c oxidase subunit 1, complex IV, mitochondrial | *Caenorhabditis elegans* | Mouse (monoclonal) | 1/1,000 | Thermo Fisher Scientific |
| CYC-1 | Cytochrome C, mitochondrial | *Caenorhabditis elegans* | Rabbit (polyclonal) | 1/1,000 | Schagger H.H. et al. (2002) |
| ISP-1 | Cytochrome b-c1 complex subunit Rieske, mitochondrial | *Caenorhabditis elegans* | Mouse (monoclonal) | 1/1,000 | Thermo Fisher Scientific |
| RISP | Rieske iron-sulfur protein | *Arabidopsis thaliana* | Rabbit (polyclonal) | 1/5,000 | Gift of Prof. Ian Small, UWA |

**Table S3.**
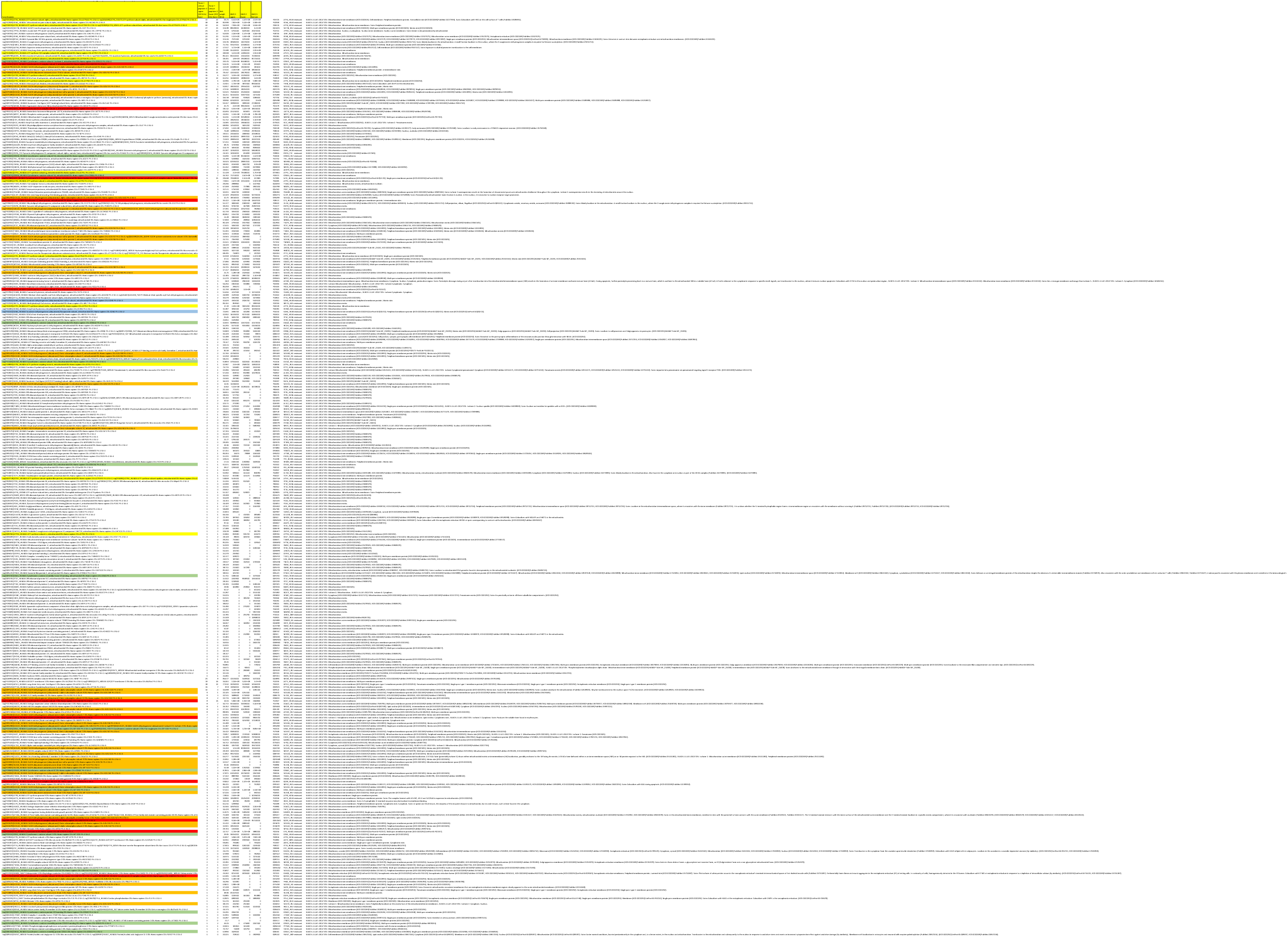
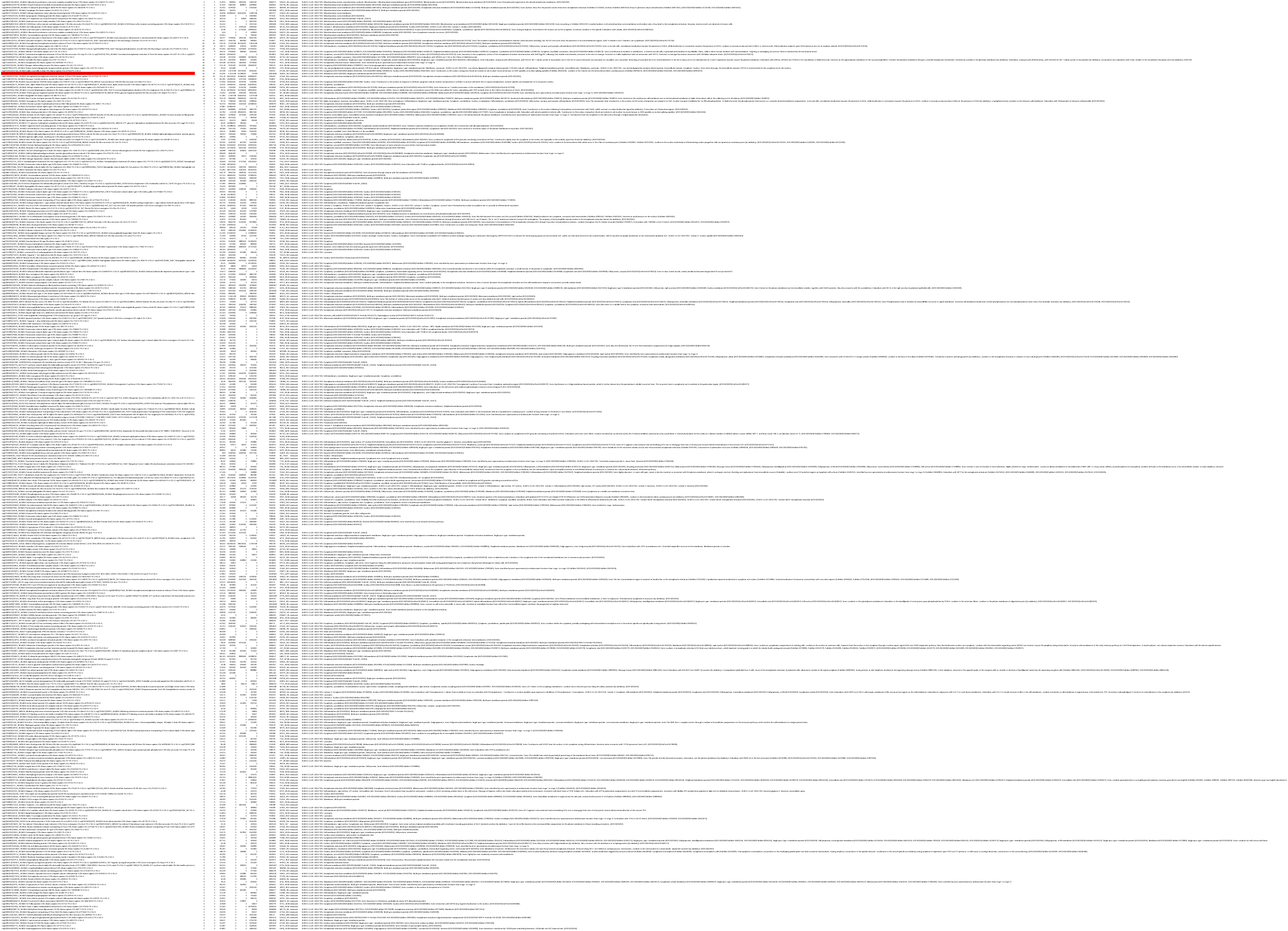
Mass-spectrometry (MS) of the ∼500 kDa (higher) and ∼350 kDa (lower) particles in human mitochondria treated with Astaxanthin. Peptides highlighted with orange color indicate to CI subunits, in blue CII subunits, in red CIII subunits, in green CIV subunits, in yellow CV (i.e., ATP-synthase).

**Table S4.**
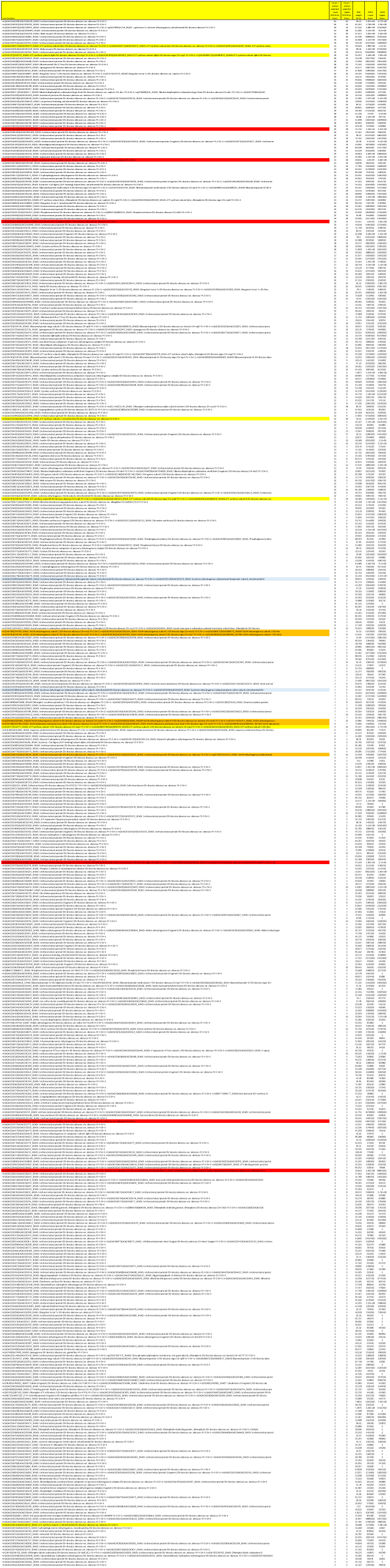
Mass-spectrometry (MS) of the ∼500 kDa (higher) and ∼350 kDa (lower) particles in cauliflower mitochondria treated with Astaxanthin. Peptides highlighted with orange color indicate to CI subunits, in blue CII subunits, in red CIII subunits, in green CIV subunits, in yellow CV (i.e., ATP-synthase).

**Figure S1.**
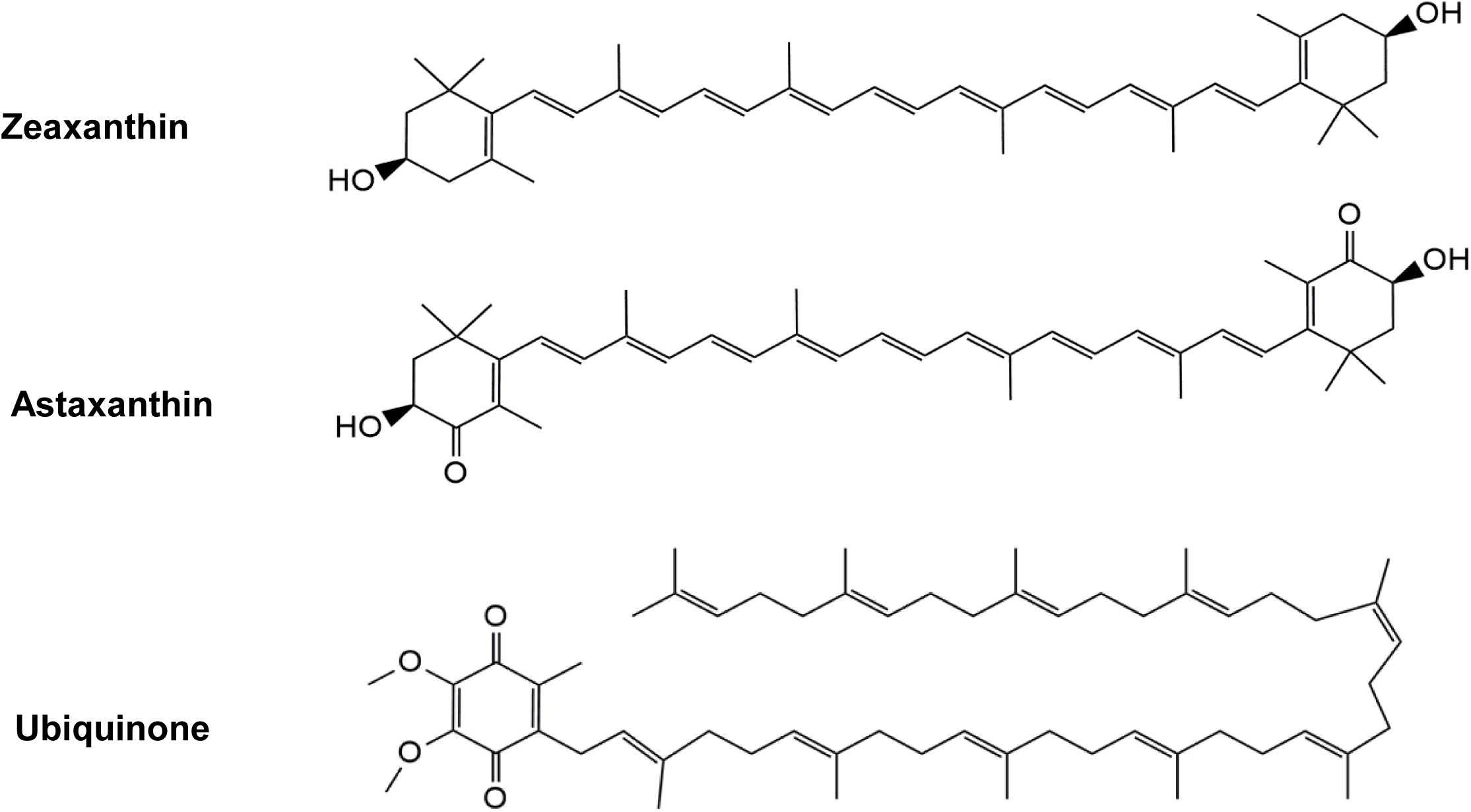
Chemical structures of the carotenoids Zeaxanthin and Astaxanthin compared with ubiquinone. Molecular structures of Zeaxanthin and Astaxanthin as well as ubiquinone.

**Figure S2.**
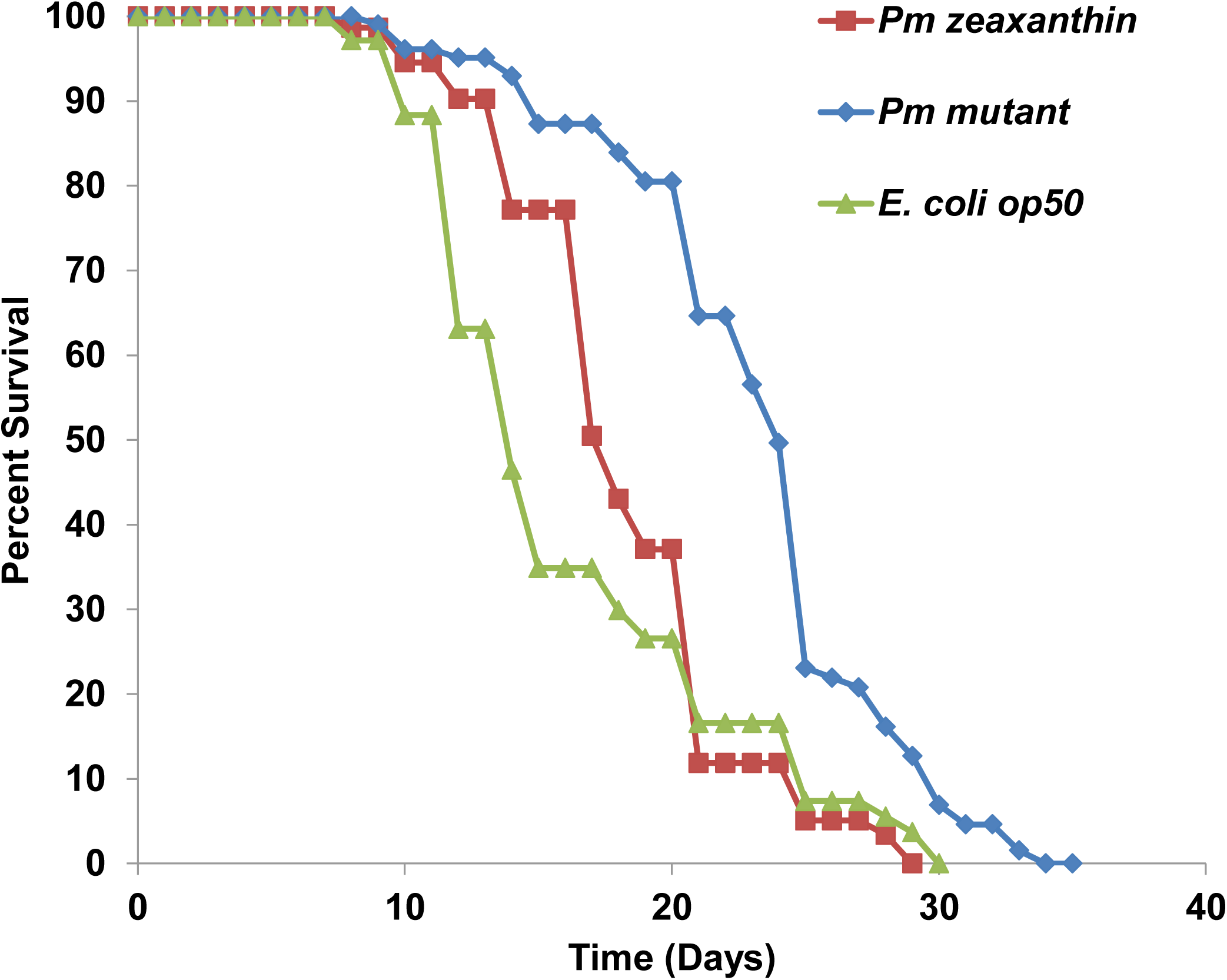
Lifespan of *C. elegans* fed with Zeaxanthin. A survival curve of nematodes supplemented with zeaxanthin in different stages of the development. About 75 animals were fed with *P. marcusii* bacteria that produce zeaxanthin (red), control *P. marcusii* bacteria (blue) or control *E. coli* OP50 bacteria (green). Animal survival was calculated with the Kaplan-Meier method, and survival differences were tested for significance using the log-rank test compared with the control. The mean lifespans (in days) of the different groups were 18.32 ± 0.53, 23.43 ± 0.57 and 16.29 ± 0.73, respectively. Lifespan curves were analyzed by plotting Kaplan-Meier survival curves (Goel*, et al.* 2010). Mean lifespan data was compared using Log-rank test (Mantel 1966) with appropriate correction for multiple comparisons OASIS-2.

**Figure S3.**
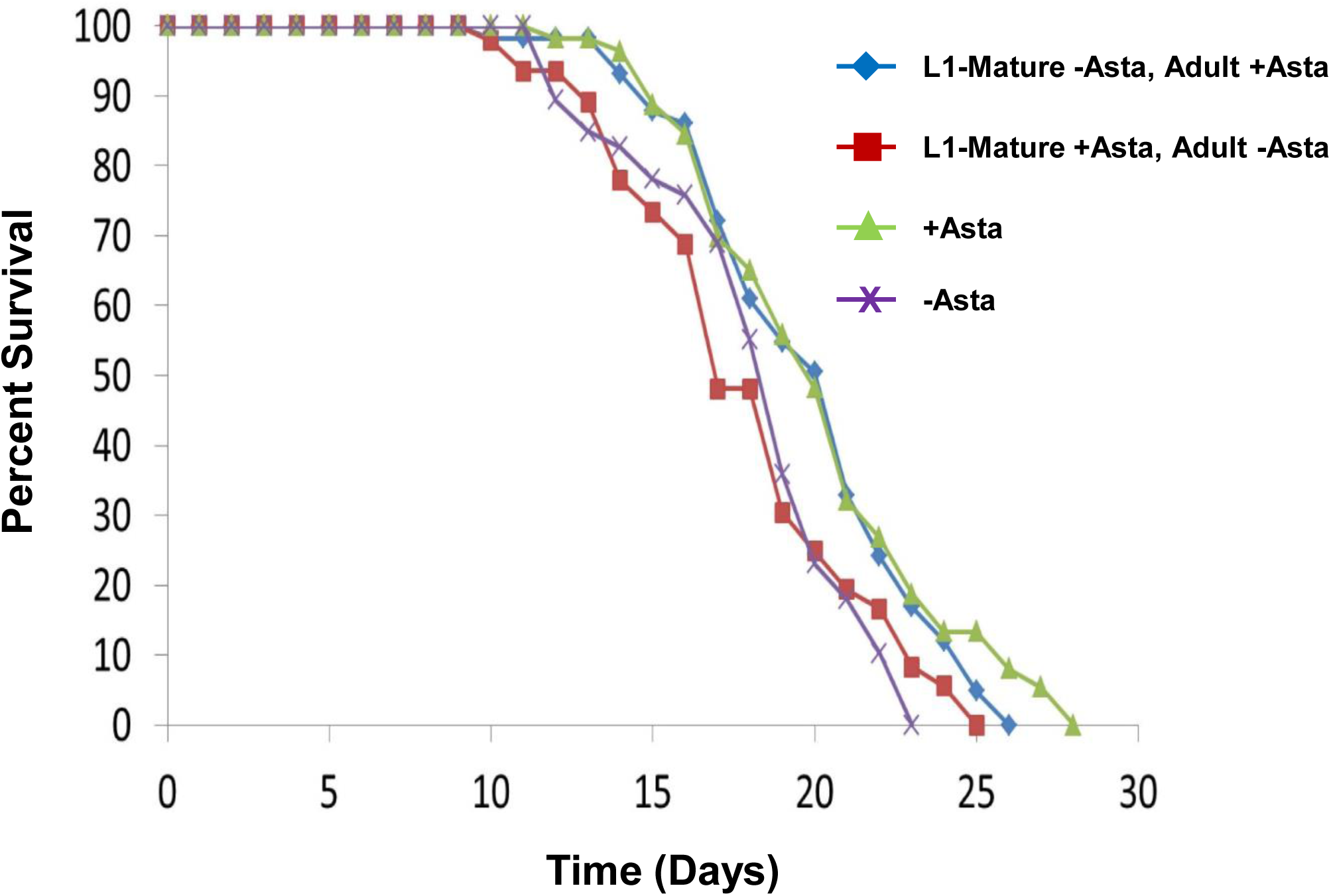
Astaxanthin effect on the median lifespan at the adult stage. A survival curve of nematodes supplemented with Astaxanthin in different stages of the development. About 75 animals were used in the experiment. Worms were fed with Paracoccus marcusii with no carotenoids expression from the embryo stage until L4 stage, and from L4 stage until their death with P. marcusii which produce Astaxanthin (blue), another group of worms were fed from the embryo stage until L4 stage with P. marcusii which produce Astaxanthin, and from L4 stage until their death with P.M. with no carotenoids expression (red). The two control groups included (i) worms which were fed from the embryo stage until their death, with *P. marcusii* with Astaxanthin (green); (ii) worms which were fed from the embryo stage until their death with *P. marcusii* with no carotenoids expression (purple). *C. elegans* survival was calculated with the Kaplan-Meier method, and survival differences were tested for significance using the log rank test compared to the control. The mean lifespans (in days) of the groups given *P. marcusii* with no carotenoids expression during development and *P. marcusii* with Astaxanthin during adulthood, *P. marcusii* with Astaxanthin during development and *P. marcusii* with no carotenoids expression during adulthood, *P. marcusii* with Astaxanthin during all their lives and *P. marcusii* with no carotenoids expression during all their lives were 19.87 ± 0.48,17.95 ±0.57, 20.18 ± 0.55 and 18.22 ± 0.49, respectively. Lifespan curves were analyzed by plotting Kaplan-Meier survival curves (Goel*, et al.* 2010). Mean lifespan data was compared using Log-rank test (Mantel 1966) with appropriate correction for multiple comparisons OASIS-2.

**Figure S4.**
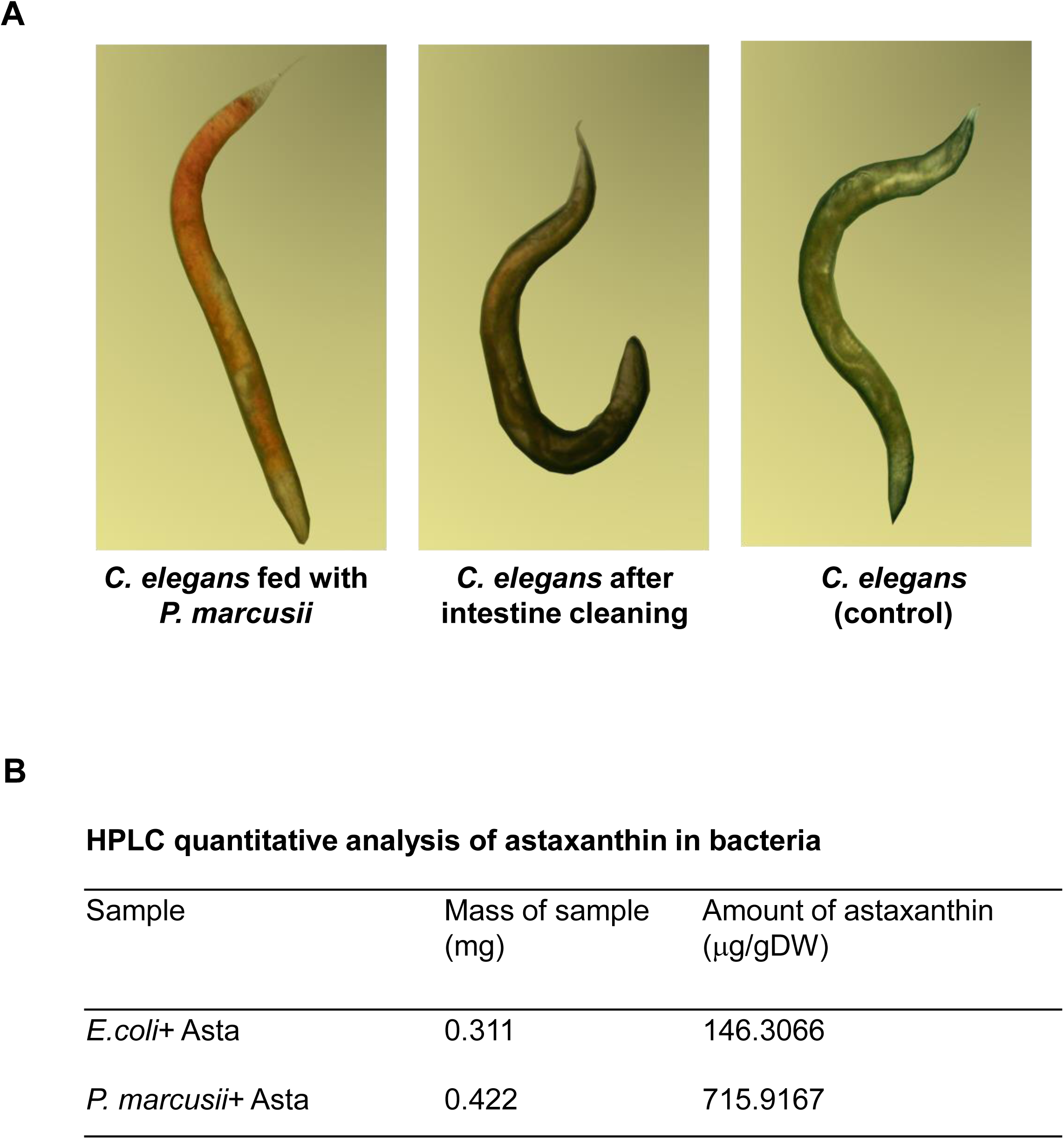
Astaxanthin from *P. marcusii* is absorbed by *C. elegans* cells. (A). Stereo microscope photograph of Astaxanthin-fed animals. *C. elegans* fed with *P. marcusii* expressing Astaxanthin from the embryo stage for 5 days (left panel). After 5 days the worms were fed with *E. coli* bacteria for 24 hours, in order to wash their intestines (middle panel). The control worms were fed just with *E. coli* bacteria (right panel). (B). Quantification of Astaxanthin in *P. marcusii* and in *E. coli* cells by HPLC. The amount of Astaxanthin is shown as microgram per gram dry weight (μg/gDW). Stereomicroscope images of Astaxanthin-fed animals. *C. elegans* were fed with *P. marcusii* containing Astaxanthin (left and middle panels) or control *P. marcusii* (left panel) from the embryo stage until adulthood. Then one group of worms were fed with *E. coli* for 24 hours, which cleans their intestine (middle panel).

**Figure S5.**
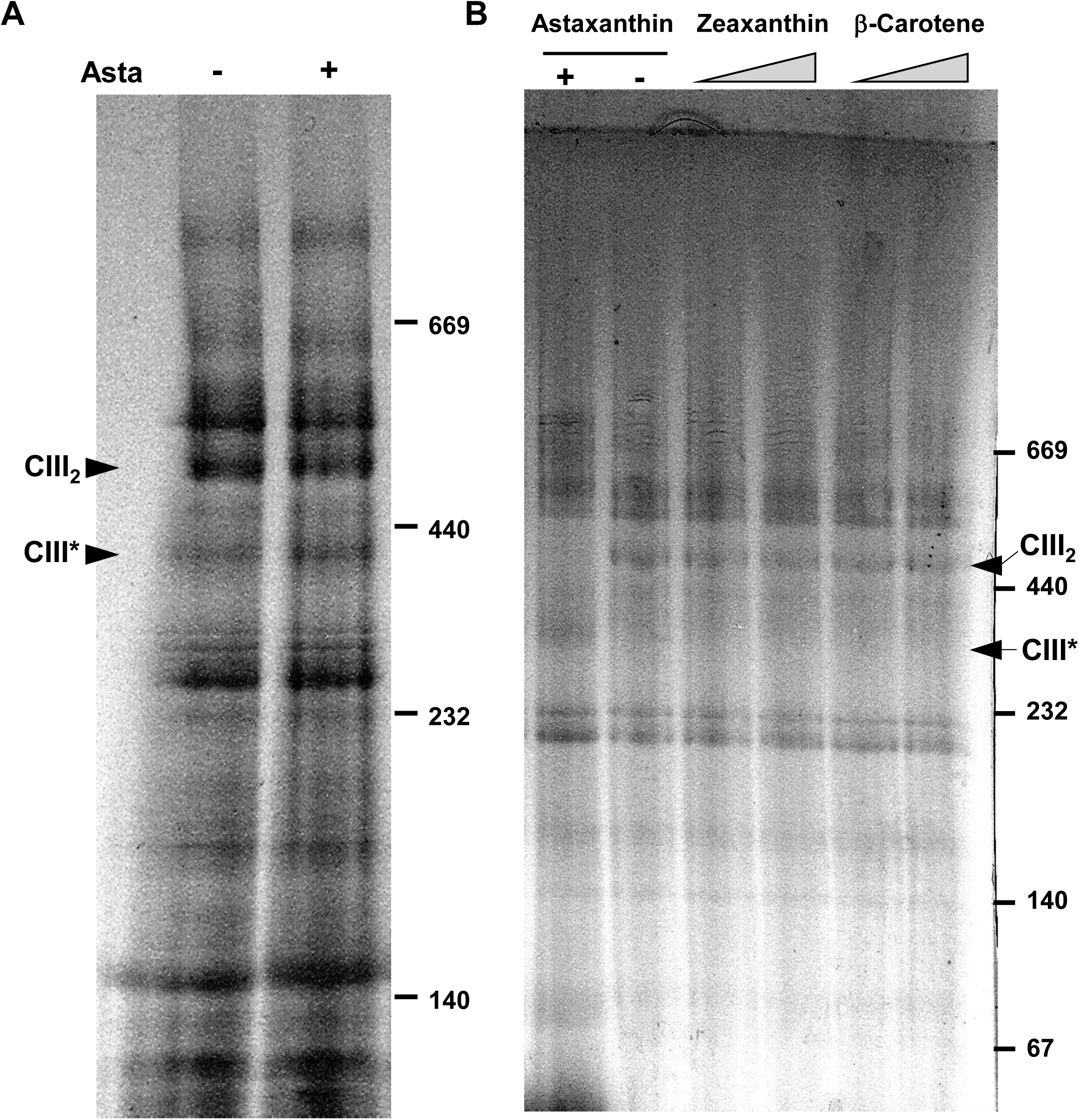
Accumulation of respiratory complexes in *C. elegans* mitochondria treated with astaxanthin. (**A**). Crude mitochondria were obtained from *C. elegans* grown in the absence (-) and presence of *P. marcusii*. Mitochondrial complexes were separated by BN-PAGE. (**B**). Crude mitochondria obtained from *C. elegans* were treated in absence (-) or presence (+) of Astaxanthin (10 μM), Lycopene (5 and 10 μM), and Zeaxanthin (5 and 10 μM), and then separated by BN-PAGE. The positions of the holo-complex III dimer (CIII_2_) and sub-CIII (CIII*) particles are indicated by arrows.

**Figure S6.**
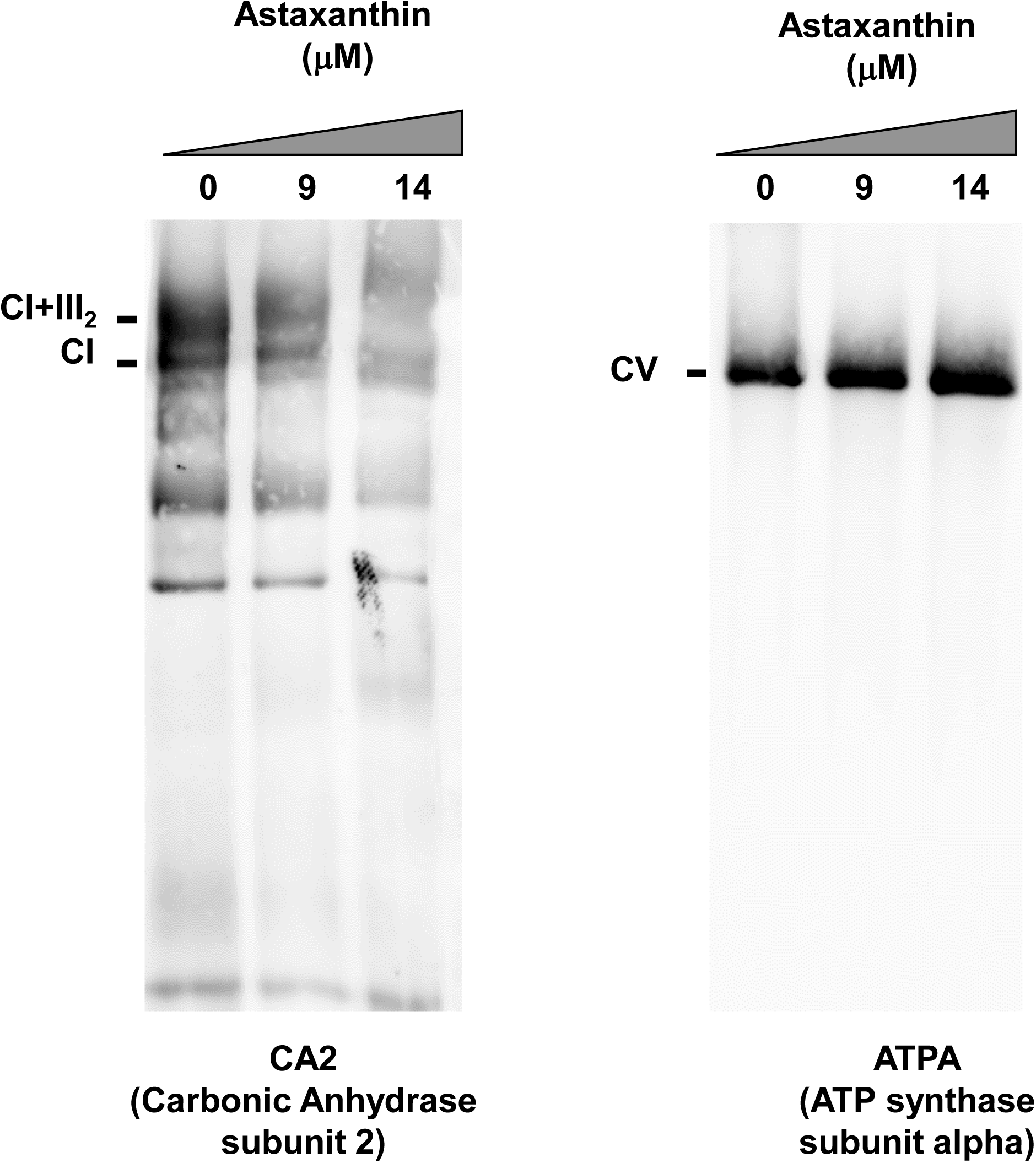
Accumulation of respiratory complexes in cauliflower mitochondria treated with different concentrations of Astaxanthin. Astaxanthin, at various concentrations (as indicated in the blots), was added to cauliflower mitochondria. The mitochondrial complexes were then separated by BN-PAGE, and the gel was stained by Coomassie brilliant blue. Antibodies raised against CA2 (Carbonic anhydrase subunit 2) and ATPA (ATP synthase subunit alpha) were used to assay the levels and integrity of the organellar complexes. The positions of the native CI (∼1,000 kDa), CV (about 600 kDa) and supercomplex I+III are indicated.

